# The Human Cytomegalovirus vGPCR UL33 is Essential for Efficient Lytic Replication in Epithelial Cells

**DOI:** 10.1101/2024.09.18.609710

**Authors:** MacKenzie R. Freeman, Abigail L. Dooley, Matthew J. Beucler, Wes Sanders, Nathaniel J. Moorman, Christine M. O’Connor, William E. Miller

**Affiliations:** Department of Molecular and Cellular Biosciences, and the Graduate Program in Molecular Genetics, Biochemistry, and Microbiology, University of Cincinnati College of Medicine, Cincinnati, OH, USA; Infection Biology, Lerner Research Institute, Cleveland Clinic, Cleveland, OH, USA; Molecular Medicine, Cleveland Clinic Lerner College of Medicine of Case Western Reserve University, Cleveland Clinic, Cleveland, OH, USA; Sheikha Fatima bint Mubarek Global Center for Pathogen and Human Health Research, Cleveland Clinic, Cleveland, OH, USA; Department of Microbiology and Immunology, University of North Carolina at Chapel Hill, Chapel Hill, NC, USA; Case Comprehensive Cancer Center, Cleveland, OH, USA

**Keywords:** HCMV, cytomegalovirus, latency, lytic replication, salivary gland, vGPCRs, UL33, Gαq Running

## Abstract

Human cytomegalovirus (HCMV) is a β-herpesvirus which is ubiquitous in the human population. HCMV has the largest genome of all known human herpesviruses, and thus encodes a large array of proteins that affect pathogenesis in different cell types. Given the large genome and the ability of HCMV to replicate in a range of cells, investigators have begun to identify viral proteins required for cell type-specific replication. There are four proteins encoded in the HCMV genome that are homologous to human G protein-coupled receptors (GPCRs); these viral-encoded GPCRs (vGPCRs) are UL33, UL78, US27, and US28. In the current study, we find that deletion of all four vGPCR genes from a clinical isolate of HCMV severely attenuates lytic replication in both primary human salivary gland epithelial cells, as well as ARPE-19 retinal epithelial cells as evidenced by significant decreases in immediate early gene expression and virus production. Deletion of UL33 from the HCMV genome also results in a failure to efficiently replicate in epithelial cells, and this defect is manifested by decreased levels of immediate early, early, and late gene expression, as well as reduced viral production. We find that similar to US28, UL33 constitutively activates Gαq-dependent PLC-β signaling to high levels in these epithelial cells. We also find that UL33 transcription is more complicated than originally believed, and there is the potential for the virus to utilize various 5’ UTRs to create novel UL33 proteins that are all capable of constitutive Gαq signaling. Taken together, these studies suggest that UL33 driven signaling is important for lytic HCMV replication in cells of epithelial origin.

## Introduction

Human cytomegalovirus (HCMV) is a β-herpesvirus which is ubiquitous in the human population, with approximately 60-90% of adults seropositive for this virus (1). HCMV is generally spread through saliva, though it is also transmitted through other bodily fluids and spread vertically from mother to fetus during pregnancy. While HCMV infection only causes mild cold- or flu-like symptoms in immunocompetent individuals, infection can cause side effects such as gastrointestinal ulceration, hepatitis, pneumonitis, and retinitis in immunocompromised patients (2). Congenital HCMV infection is the most common congenital infection in the United States; approximately 10-15% of infected newborns will suffer from side effects such as encephalitis, microcephaly, hepatitis/hepatomegaly, nephritis, and colitis (3, 4). Additionally, congenital HCMV infection is the leading viral cause of neurodevelopmental delay, as well as the leading non-genetic cause of sensorineural hearing loss in children (5–8).

A distinct characteristic unique to herpesviruses is their ability to go through multiple stages of infection within a host; these stages include lytic replication, latency, and reactivation (9). Initial infection is typically lytic and occurs in the nasopharyngeal epithelium, while latency occurs in hematopoietic progenitor cells and other cells of the myeloid lineage. Following primary infection or following lytic reactivation from latency, the virus can persistently replicate in the salivary epithelium where it can contribute substantially to horizontal spread (10). During lytic infection, viral replication is separated into three stages: immediate early, early, and late (11). While lytic replication most commonly occurs in epithelial cells *in vivo*, lytic infections can also be modeled in many different cell types *in vitro*, such as fibroblast, epithelial, and endothelial cells (12–15). Defining the viral mechanisms and genes that govern cell-specific replication remains an important area of investigation.

G protein-coupled receptors (GPCRs) are cell surface proteins with seven transmembrane domains. The amino (N) terminus of the protein extends into the extracellular space, while the carboxy (C) terminus extends into the cytoplasm (16). GPCRs can trigger signaling cascades by associating with heterotrimeric G proteins, which are made up of alpha (Gα), beta (Gβ), and gamma (Gγ) subunits (17, 18). The HCMV genome encodes four proteins with homology to GPCRs; these are UL33, UL78, US27, and US28 (19–22). These viral GPCRs (vGPCRs) are also found in the viral envelope, suggesting they may function in initial aspects of infection, such as facilitating interactions with target host cells or driving immediate signaling that is a positive factor in the establishment of lytic or latent infections (23–27). While the vGPCRs are well-conserved and present in all sequenced viral isolates, they are largely dispensable for lytic replication in fibroblasts *in vitro*. HCMV US28, the most well-characterized of the vGPCRs, signals through multiple G-proteins and has demonstrable roles in latency and cellular migration (24, 28–30). Much less is known about the other three vGPCRs, and moreover, the strict species-specificity of HCMV has significantly impeded the ability to conduct *in vivo* studies investigating their function. However, the HCMV UL33 and UL78 vGPCRs have orthologs in rodent CMVs, which allows investigations into potential roles for these orthologs in *in vivo* settings. UL33 orthologs in rodent CMVs are required for proper replication and pathogenesis *in vivo*. The murine CMV UL33 ortholog, M33, is required both for efficient dissemination to the salivary gland, as well as replication once the virus reaches the salivary gland (31–33). The rat CMV UL33 ortholog, R33, also appears to play a major role in pathogenesis, as the deletion of R33 from RCMV causes higher survival rates and decreased replication within the salivary glands (34). Taken together, these data suggest that rodent CMV orthologs of HCMV UL33 play an important role in viral replication and pathogenesis, particularly in the salivary epithelium. However, the role of UL33 in HCMV pathogenesis remains largely unclear.

Though UL33 is characterized as a putative chemokine receptor within the GPCR superfamily, it remains an ‘orphan receptor’, as its ability to bind to or respond to chemokines is unknown. UL33 has some ability to nominally activate G protein signaling cascades, although this activity remains controversial and differs from study to study (35–37). UL33 is also unique among the four HCMV vGPCRs, as the gene is comprised of multiple exons which are spliced to create the full-length UL33 mRNA. Interestingly, these exons contain multiple methionine start sites, indicating this gene could give rise to several unique proteins. Additionally, UL33 enhances *in vitro* virus dissemination through both extracellular and cell-to-cell routes of infection in fibroblasts (38), functions during reactivation from latency by modulating the activity of the viral major immediate early promoter (MIEP) (39), and induces trophoblast migration that could contribute to congenital transmission (37). Moreover, UL33 modulates CXCL12-induced THP-1 monocytic cell migration (40). CXCL12 signaling is also modulated by US27 upon its colocalization with CXCR4 in HEK and fibroblast cells (41). Since the CXCL12-CXCR4 signaling axis is important in hematopoiesis, such pathway cross-talk may function to hone latently-infected cells to the bone marrow (40, 41), though US27 is not expressed during latency (24, 42). While UL33 does display activity in HCMV infected fibroblasts, hematopoietic cells, and trophoblasts, it remains unclear whether UL33 has additional roles in viral pathogenesis in other biologically-relevant cells. In particular, it remains unknown if HCMV UL33 promotes viral pathogenesis and replication in cells of epithelial origin, such as those comprising the epithelium of the salivary gland.

In this study, we characterize UL33 activity in epithelial cells to gain a new perspective regarding the biological functions of these vGPCRs during viral replication and pathogenesis, with a goal of better understanding how UL33 and the other vGPCRs may facilitate HCMV replication and persistence in the salivary epithelium.

## Results

### Deletion of vGPCR genes from the HCMV TB40/E genome causes decreased viral infectivity and replication in salivary-derived epithelial cells

Since HCMV dissemination through the human population is closely tied to HCMV virions present in saliva and urine (3, 10), we were interested in determining the biological mechanisms by which the vGPCRs contribute to HCMV infection and replication in salivary epithelial cells. Our lab has recently developed a primary salivary epithelial cell model that has proven to be a useful *in vitro* tool for the study of HCMV replication in the salivary epithelium (13). Parotid or submandibular salivary gland tissues from de-identified human subjects were used to establish primary human salivary gland epithelial (hSGE) cell lines. These hSGE lines were characterized in detail and demonstrate many characteristics of salivary acinar epithelial cells including expression of acinar specific genes such as amylase, histatin-1, statherin, etc. (13). When grown either as three-dimensional “salispheres” on Matrigel or as epithelial monolayers on plastic cell culture dishes, we can efficiently infect these hSGEs with various epitheliotropic strains of HCMV, including TB40/E, which initiates viral gene expression and produces infectious virions after infection of these cells (13, 43). HCMV infection of and replication within epithelial cells is not as efficient as in fibroblasts, due to additional viral genes and processes necessary for replication in epithelial cells (13, 43, 44). When compared to the ability of TB40/E to infect HS68 primary human fibroblasts [as assessed by IE1 (immediate early gene 1) gene expression at 48 hours post-infection (hpi)], we generate what we term the “hSGE:HS68 infectivity index.”

To determine what effect the vGPCRs have on the hSGE:HS68 infectivity index, we infected hSGE cells and HS68 fibroblasts (MOI= 2 IU/cell) with either wild type HCMV, TB40/E*mCherry* (WT) (45) or a variant deleted for all four vGPCR genes, TB40/E*mCherry*-allΔ (Δall) (46), and assessed immediate early protein (IE1/IE2) expression at 48 hpi. We find that TB40/E-WT infection generates an infectivity index of 0.32, while the TB40/E-Δall vGPCR mutant generates an infectivity index of 0.068 (**Fig. 1A**), indicating the Δall vGPCR mutant is 78% defective at initiating IE gene expression in hSGE cells compared to WT virus. We then analyzed viral production with WT and Δall viruses by harvesting cell-free virus from infected hSGE cells at time points between 2- and 32-days post-infection (dpi). Viral titers in the hSGE culture media were determined using a flow-based assay as described previously (13). While the TB40/E-WT virus replicated and grew to a titer of approximately 5×10^4^ IU/mL, the TB40/E-Δall vGPCR mutant failed to replicate (**Fig. 1B**). We conclude that the hSGEs infected with TB40/E-Δall results in fewer infectious particles compared to TB40/E-WT, consistent with our findings regarding the inability of the TB40/E-Δall vGPCR mutant to efficiently initiate IE gene expression in the hSGE cells (**Figs. 1A, B**). Given that the vGPCR null mutant exhibits a defect in initiating IE gene expression and efficient replication in the hSGE cells, we next determined if this defect is recapitulated in the immortalized ARPE-19 retinal pigment epithelial cell line, a cell line commonly used to study HCMV infection of epithelial cells. We therefore infected ARPE-19 cells with TB40/E-WT or -Δall (MOI= 2 IU/ml) and calculated ARPE:HS68 infectivity indices. TB40/E-WT infection generates an infectivity index of 0.17, while TB40/E-Δall generates an infectivity index of 0.02 (**Fig. 1C**), indicating the Δall vGPCR mutant is 88% defective at initiating IE gene expression in ARPE-19 cells compared to WT virus. Therefore, although the WT virus infectivity index is lower in ARPE-19 cells than what we observed in the hSGE cells, the ARPE-19 cells display a similar trend. Thus, in subsequent experiments in this study, we will use ARPE-19 cells to probe the biological and biochemical functions of the largely uncharacterized HCMV vGPCR, UL33.

**Figure 1.**
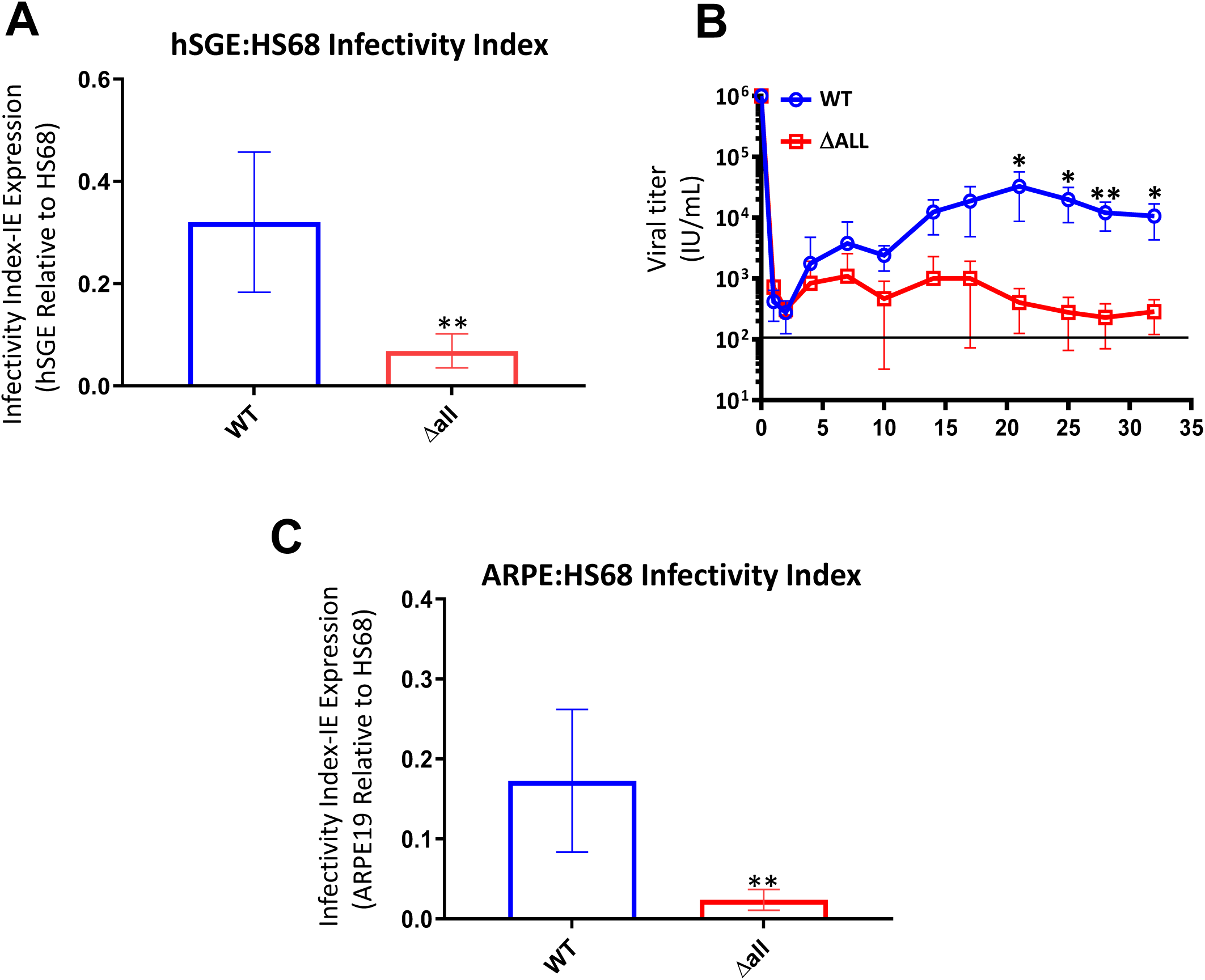
Infection and replication of a HCMV mutant deleted for all four vGPCRs. (**A**) Primary human salivary gland epithelial cells (hSGEs) were infected (MOI= 2.0 IU/cell) with TB40/E-*mCherry* or TB40/E-mCherry(Δall). Concurrently, equal numbers of HS68 HFF cells were infected with the same amount of virus. At 2 days post-infection (dpi), cells were analyzed by flow cytometry for the proportion of cells which were IE1/2 positive. Infectivity Indices were calculated by taking the percent of IE1/2 positive hSGE cells and dividing by the percent IE1/2 positive HS68 cells from the same experiment. The results are representative of 8 independent experiments with 2 different parotid and 2 different submandibular cell cultures. Error is shown as Standard deviation and significance was calculated using unpaired t-tests. (**B**) Primary hSGE cells derived from 2 different parotid glands and 2 different submandibular glands and were infected (2.0 IU/cell) and supernatants harvested at the indicated dpi. Supernatants were used to calculate viral titers on HS68 HFF cells as described in the materials and methods. The results are representative of 4 independent experiments. Error is shown as Standard deviation and significance was calculated using unpaired t-tests. **(C)** ARPE-19 or HS68 HFF cells were infected as in panel A and Infectivity Indexes were calculated by taking the percent of IE1/IE2 positive ARPE cells and dividing by the percent of IE1/IE2 positive HS68 cells. The results are representative of 10 independent experiments. Error is shown as Standard deviation and significance was calculated using unpaired t-tests. \**p*<0.05, \*\**p*<0.01

### The four HCMV-encoded vGPCRs, including UL33, are expressed in infected ARPE-19 epithelial cells

Given that the HCMV vGPCRs play a critical role in promoting viral replication in hSGE and ARPE-19 cells, we next evaluated the expression and kinetics of each vGPCR following infection of ARPE-19 cells. Our prior work and that of others revealed the vGPCRs are incorporated into the mature viral particle (24, 26, 27, 45, 47), and thus could function in processes immediately following virion entry. This could include the initiation of IE gene expression, as we note in **Figs. 1A, C**. Further, while we and others have shown UL78 (27) and US28 (48, 49) are each expressed during epithelial infection, a comparison of the expression and kinetics for all four vGPCRs during epithelial infection, as well as how they contribute to sustained viral production, as in **Fig. 1B**, remains incomplete. To examine vGPCR protein abundance in ARPE-19 cells, we used four different, previously characterized TB40/E recombinants, each generated in the TB40/E*mCherry* background. Each recombinant contains a triple-FLAG epitope tag on the carboxy (C) terminus of one of the vGPCR genes: TB40/E*mCherry*-UL33-3xF (UL33-3xF) (39), TB40/E*mCherry*-UL78-3xF (UL78-3xF) (27), TB40/E*mCherry*-US27-3xF (US27-3xF) (45), TB40/E*mCherry*-US28-3xF (US28-3xF) (46). We placed the epitope tag on the C-terminus, as previous studies with several different GPCRs have revealed the C-terminal tag does not affect the subcellular localization or signaling activity of the GPCR proteins (50–52). To ensure a high percentage of infection, we infected ARPE-19 cells (MOI= 10 IU/ml) and collected whole-cell protein lysates 48 and 96 hpi. Each vGPCR protein was then immunoprecipitated using M2-agarose beads to pull down the FLAG-tagged vGPCR, which were then separated by SDS-PAGE and subjected to western blotting with anti-FLAG HRP-conjugated antibody. TB40/E-WT contains no FLAG-tagged proteins and thus serves as the negative control for these studies (**Fig. 2A**). We find that UL33 and US27 exhibit delayed early or late kinetics with undetectable expression at 48 hpi, but increased abundance at 96 hpi (**Figs. 2B, D**). Interestingly, we find that UL78 exhibits early kinetics with robust expression at both 48 and 96 hpi (**Fig. 2C**). Finally, US28 exhibits modest expression at the early 48-hour time point, with increased expression at the later 96-hour time point (**Fig. 2E**). This correlates to previously published data, which suggested that UL78 and US28 are expressed with early kinetics, and UL33 and US27 are expressed with late kinetics (23, 25–27, 39). We note that GPCRs are notorious for complicated banding patterns on western blots due to post-translational modifications, such as glycosylation, as well as the tendency of these proteins to aggregate in detergent cell extracts. This results in smearing, as well as the appearance of apparent dimers, consistent with our current (**Fig. 2**) and prior observations (27, 39, 45, 46, 49, 51, 53, 54).

**Figure 2.**
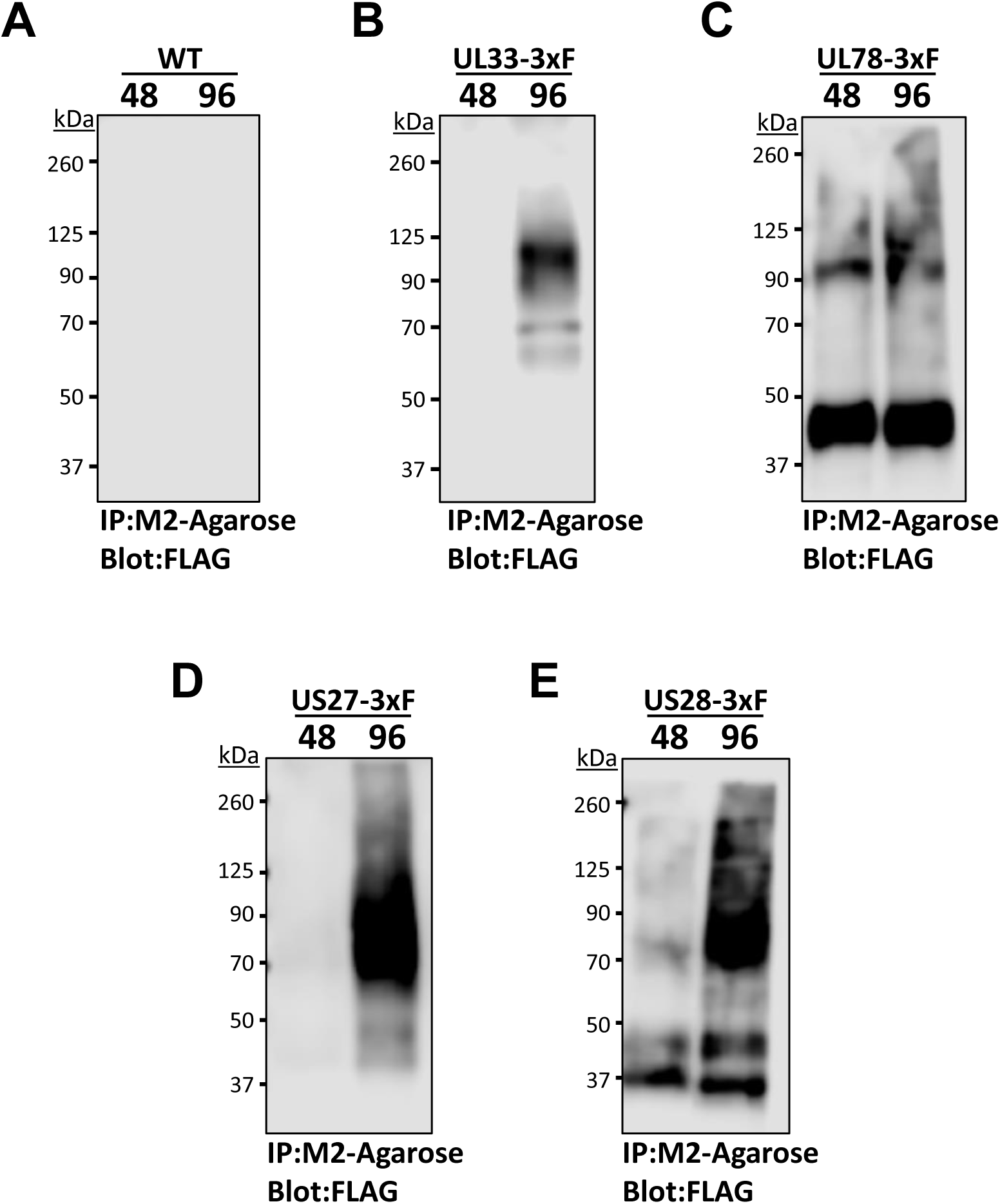
The four HCMV vGPCR proteins are expressed during TB40/E infection of ARPE-19 cells. ARPE-19 cells were infected (MOI= 10.0 IU/cell) with various TB40/E-based viruses: (**A**) WT, (**B**) UL33-3xF, (**C**) UL78-3xF, (**D**) US27-3xF, or (**E**) US28-3xF. Protein lysates were harvested at either 48 or 96 hours post-infection (hpi) and immunoprecipitated utilizing anti-FLAG M2 agarose beads. Samples were then probed for M2 (FLAG) to detect the abundance of each vGPCR. The western blots shown are representative of at least three independent experiments.

### HCMV UL33 displays late expression kinetics and is localized to the viral assembly complex

The impact of UL33 expression during epithelial cell infection remains elusive, and thus, we focused on deciphering the contribution of UL33 to infection in this cell type. To this end, we generated two, independent UL33 deletion mutants using UL33-3xF as the parental strain. Using galK-mediated recombineering (55), we deleted the entire open reading frame (ORF), inclusive of the triple FLAG tag, to generate the UL33 deletion mutants termed TB40/E*mCherry*-UL33Δ1 (UL33Δ1) and UL33Δ2 (39). Indeed, deletion of the ORF results in ablation of UL33 expression by both immunoblot (**Fig. 3A**) and immunofluorescence (**Fig. 3B**). Importantly, the UL33 deletion mutant viruses display wild type growth kinetics during lytic infection of highly permissive fibroblasts (39). Furthermore, we evaluated UL33 localization and expression kinetics in lytically-infected fibroblasts. Our data reveal UL33 localizes to the perinuclear, viral assembly complex (VAC; **Fig. 4A**), a virus-specific structure that forms late in infection as a hub for proteins involved in particle assembly and egress (56). Perhaps UL33’s localization to the VAC is unsurprising, as the other CMV-encoded GPCRs also display VAC localization (27, 45, 48). UL33, however, also displays localization to additional cytoplasmic compartments, which occurs as early as 48 hpi, persisting through 96 hpi (**Fig. 4A**). We also assessed UL33’s expression kinetics by immunoblot. Additionally, to determine the kinetic class to which UL33 belongs, we treated a parallel culture with phosphonoacetic acid (PAA). We found UL33 is sensitive to PAA treatment (**Fig. 4B**), indicating that UL33 is expressed with true late viral kinetics. Our western analysis also parallels our immunofluorescence data, which collectively reveals that UL33 expression begins at 48 hpi, with expression increasing through 96 hpi.

**Figure 3.**
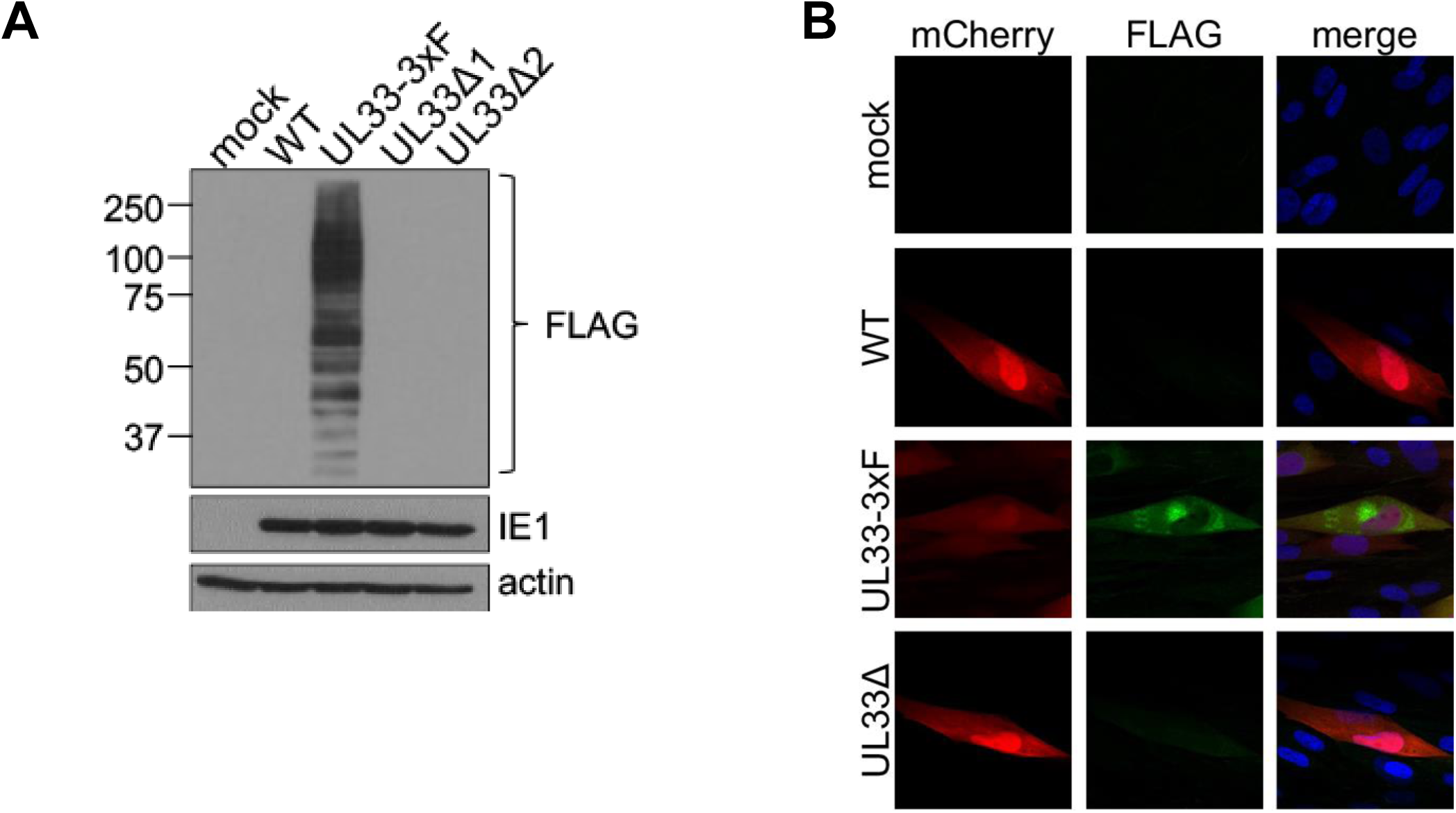
UL33 expression is ablated in TB40/E*mCherry*-UL33Δ. Fibroblasts (NuFF-1) were infected (MOI= 0.5 TCID50/cell) with WT, UL33-3xF, UL33Δ1, or UL33Δ2 and analyzed at 96 hpi. **(A)** Whole cell lysates were probed by immunoblot with the indicated antibodies. IE1, marker of lytic infection; actin, loading control. N = 3, representative blots shown. **(B)** Cells were fixed, permeabilized, and UL33 localization was assessed using an antibody against the FLAG epitopes (green). mCherry (red) is shown as a marker of infection, and DAPI (blue) was used to visualize nuclei. Images were acquired at 63X magnification. N= 3, representative images shown.

**Figure 4.**
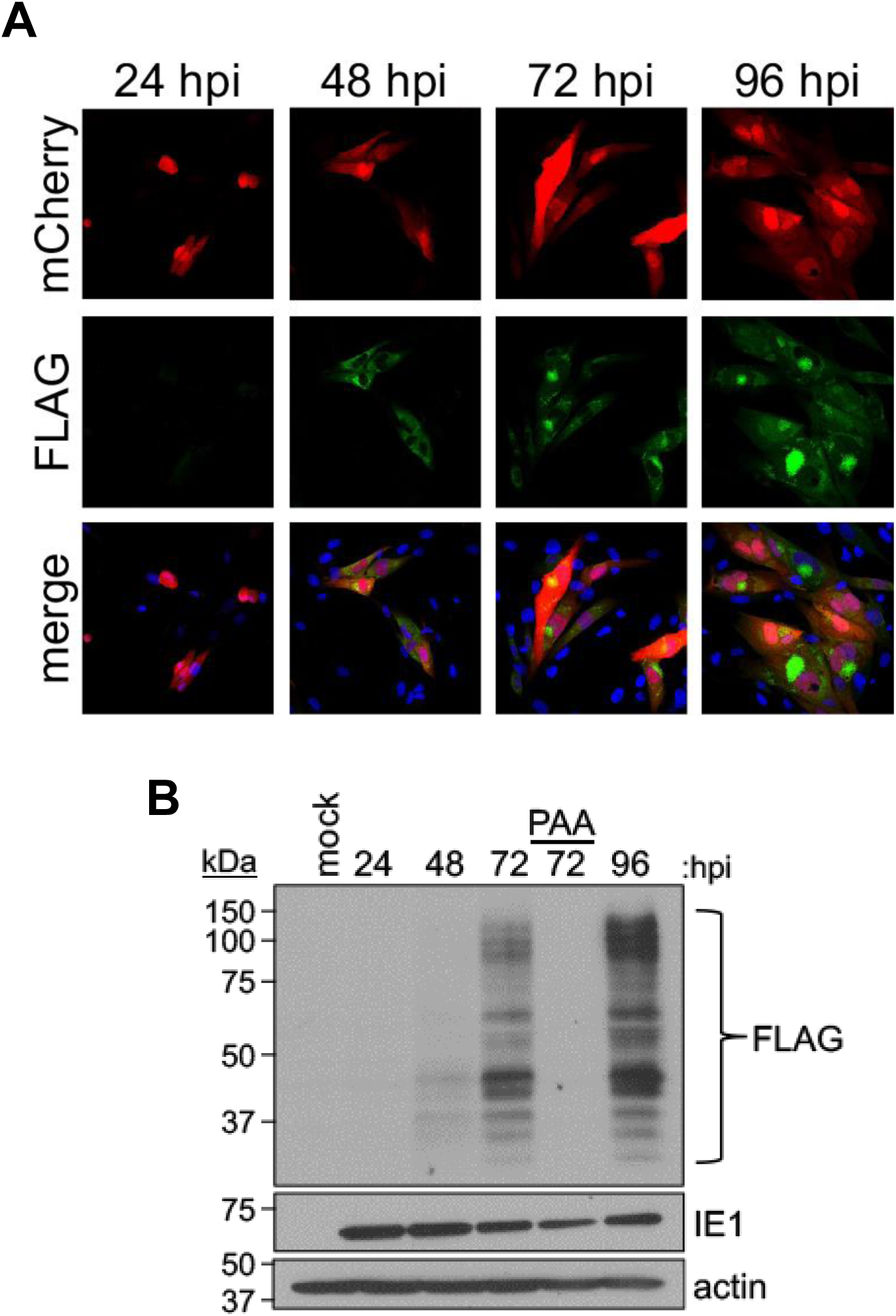
UL33 is a late protein. **(A)** Fibroblasts (NuFF-1) were infected (MOI= 0.5 TCID50/cell) with TB40/E-*mCherry*-UL33-3xF (UL33-3xF). Cells were next fixed and permeabilized over a 96 h time course, and UL33 was visualized via its FLAG epitopes (green). mCherry (red) is a marker of infection, and DAPI (blue) was used to visualize host cell nuclei. Images were acquired at 40X magnification. N = 3, representative images shown **(B)** Cells were infected as in **(A)**, with an additional, parallel culture treated with PAA post-adsorption, which was collected at 72 hpi. Cells were otherwise collected over a 96 h time course, and whole-cell lysates were immunoblotted for the FLAG epitope tag to detect pUL33-3xF. IE1 is shown as a marker of lytic infection, and actin is shown as a loading control. M, mock-infected cells, harvested at 96 hpi. N = 3, representative blots shown.

### UL33 is required for efficient viral replication and gene expression in epithelial cells

Our previous work revealed UL33 is dispensable for replication in lytically-infected fibroblasts (39), but how UL33 contributes to viral growth in epithelial cells remained unknown. Thus, we evaluated viral replication in the absence of UL33 in infected ARPE-19 cells. Since we previously characterized the UL33 mutant viruses and found that both mutants displayed similar phenotypes (39), we used one mutant (UL33Δ1, designated subsequently as UL33Δ) for the remainder of the experiments. To this end, we infected ARPE-19 cells with WT or UL33Δ at a low moi (0.1 TCID50/cell) and harvested cell-associated virus over a 30-day time course of infection. Our data reveal deletion of UL33 attenuates viral replication compared to the WT infection (**Fig. 5**), indicating UL33 is required for efficient viral replication in ARPE-19 cells.

**Figure 5.**
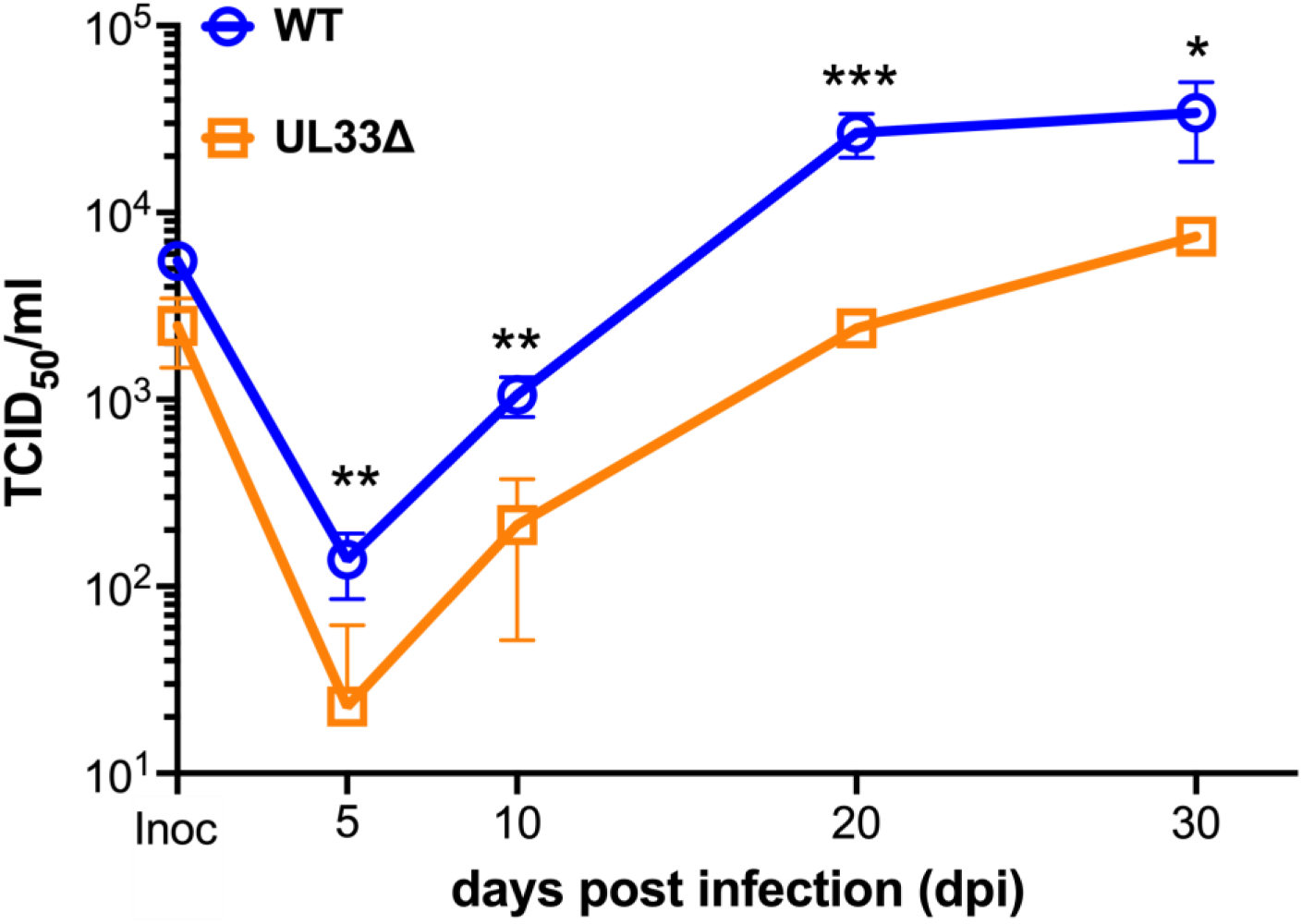
Deletion of the UL33 ORF impairs viral replication in epithelial cells. Multistep growth analyses were performed in epithelial cells (ARPE-19 cells) infected (MOI= 0.1 TCID50/cell) with the indicated viruses over a 30-day time course. Infected cells were collected at the indicated times, and viral titers of cell-associated virus were quantified by TCID50 using naïve NuFF-1 fibroblasts. N = 3 biological replicates; graph depicts one biological replicate, with 3 technical replicates per time point. Error is shown as Standard deviation and significance was calculated using unpaired t-tests. \**p*<0.05, \*\**p*<0.01, \*\*\**p*<0.005

We next asked at which stage UL33 may play a role during infection of ARPE-19 cells. To begin to address this, we evaluated protein expression from a representative protein in each kinetic class of expression: immediate early (IE), early (E), and late (L). To this end, we infected NuFF-1 fibroblasts or ARPE-19 epithelial cells with either WT or UL33Δ and collected total cell lysates over a time course of infection. As HCMV infection in ARPE-19 cells versus fibroblasts is delayed, we extended the time course by 48 h in ARPE-19 cells, as we have previously (27). As expected, UL33Δ-infected fibroblasts resulted in IE1 (IE), pUL44 (E), and pp28 (L) protein expression similar to that in WT-infected counterpart cultures (**Fig. 6A**). However, we found UL33Δ-infected ARPE-19 cells display significantly impaired IE, E, and L protein synthesis when compared to WT-infected ARPE-19 cells (**Fig. 6B**). We next asked if this phenotype we observed in the ARPE-19 cells was due to a defect in viral gene transcription. We infected ARPE-19 cells as before, and we evaluated transcription of *UL123*, *UL44*, and *UL99* (the genes encoding IE1, pUL44, and pp28, respectively). We found UL33Δ-infected ARPE-19 cells displayed a significant impairment in the transcription of all three genes (**Fig. 7**). Collectively, these findings suggest UL33 contributes to efficient viral replication in ARPE-19 epithelial cells, at a stage prior to viral RNA synthesis.

**Figure 6.**
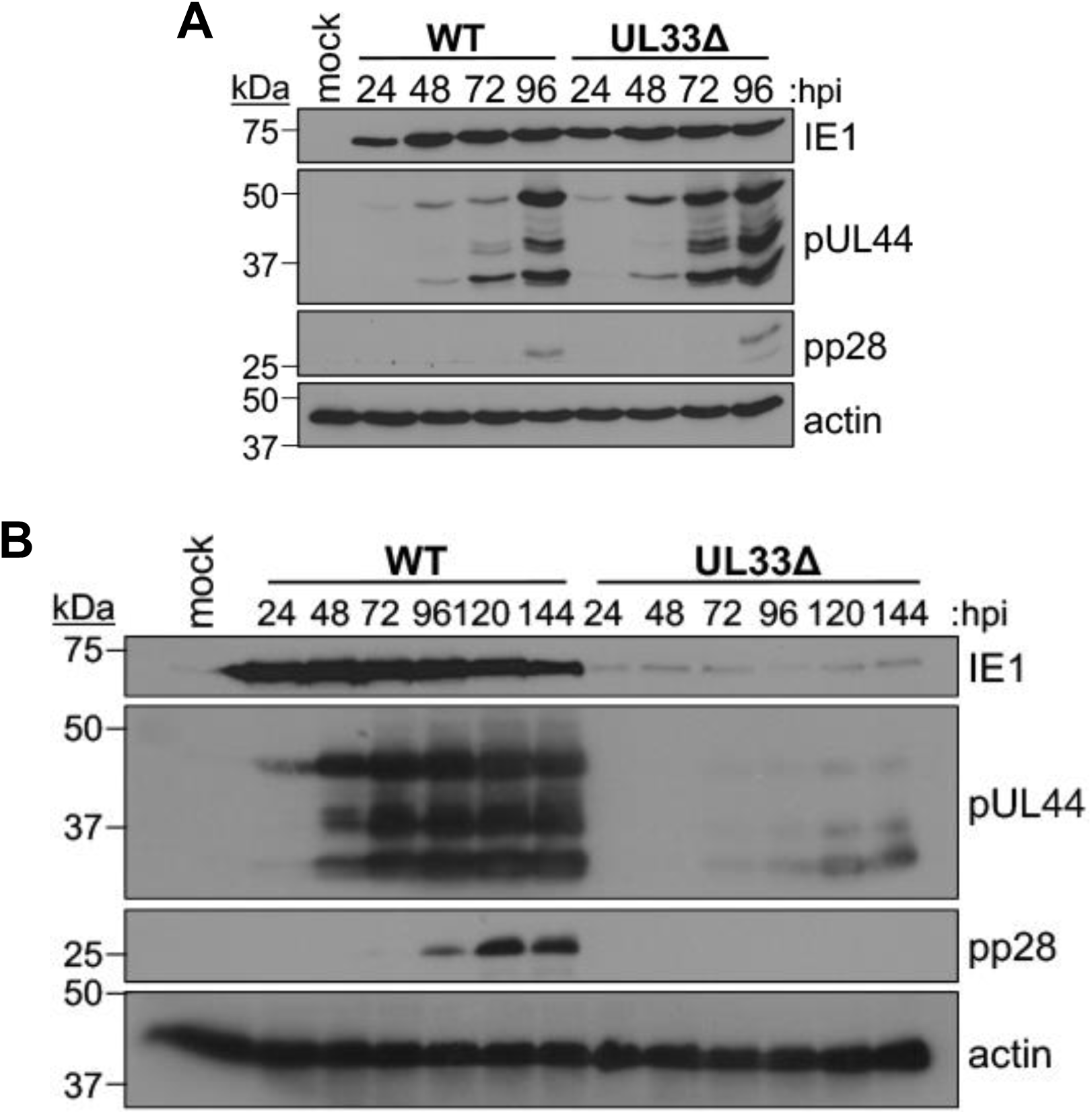
UL33 is dispensable for lytic viral protein expression in fibroblasts, but necessary in epithelial cells. **(A)** Fibroblasts (NuFF-1) or **(B)** epithelial cells (ARPE-19) were mock-, WT-, or UL33Δ-infected (MOI= 0.5 TCID50/cell) for the times indicated. Whole cell lysates were prepared and representative viral proteins from the IE (IE1), E (pUL44), and L (pp28) kinetic classes were assessed by immunoblot. Actin was assayed as a loading control. N=3, representative blots shown.

**Figure 7.**
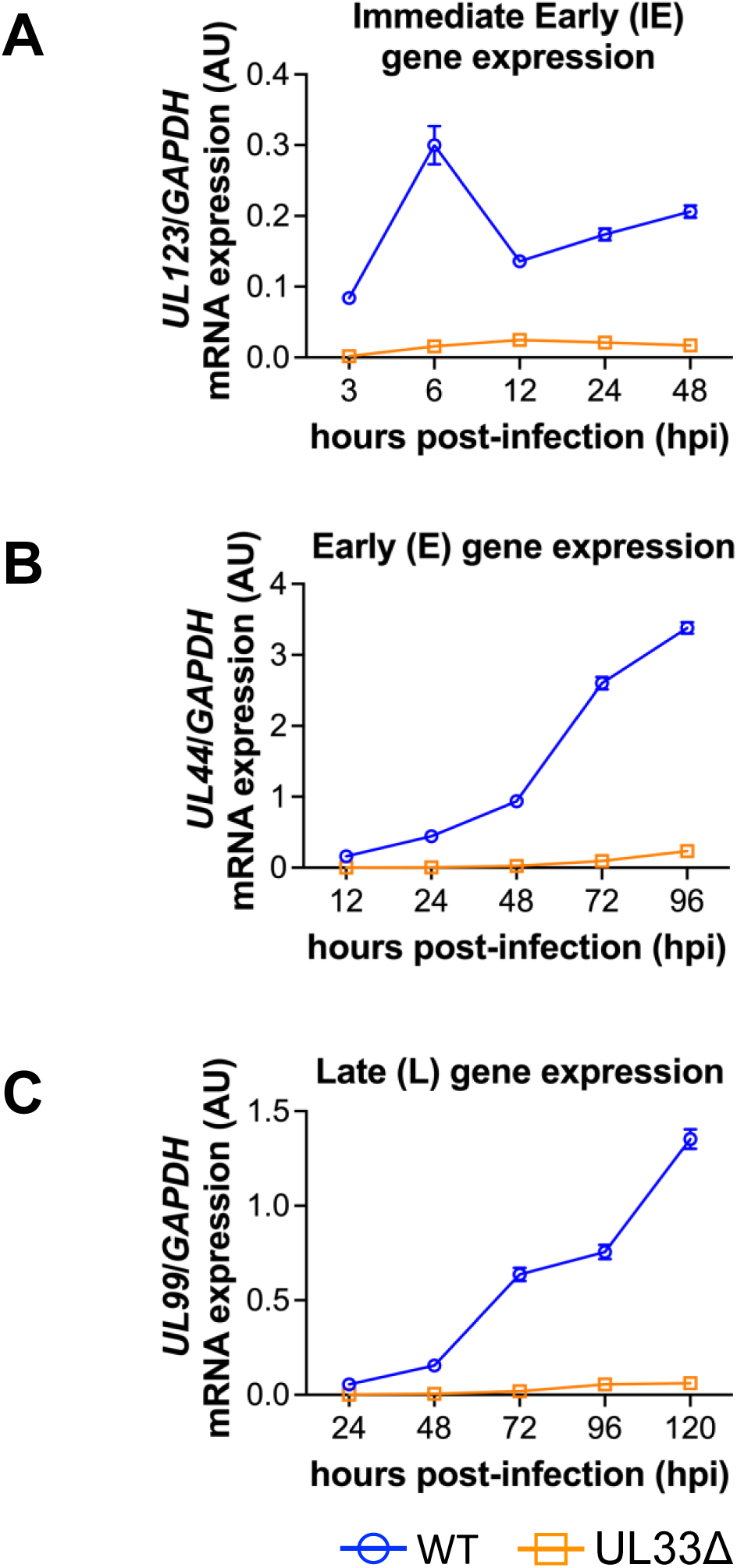
UL33 is required for efficient viral mRNA accumulation in epithelial cells. Epithelial cells (ARPE-19) were infected (MOI= 0.5 TCID50/cell) with WT (blue) or UL33Δ (orange) for the times indicated. RNA was isolated and RT-qPCR performed with primers directed at **(A)** *UL123* (IE gene), **(B)** *UL44* (E gene), and **(C)** *UL99* (L gene). **(A-C)** All samples were analyzed in technical triplicate and normalized to cellular *GAPDH*. N=4 biological replicates; representative biological replicate shown for each gene (each with 3 technical replicates). Error is shown as standard deviation, and significance was calculated using unpaired t-test. Each time point for all genes showed statistical significance of *p*<0.0001 between WT and UL33Δ.

### Both US28 and UL33N signal through the Gαq G-protein, leading to increased PLC-β activity in epithelial cells

Our data indicate that UL33 contributes to the successful infection of epithelial cells, however the underlying mechanism is unclear. As a GPCR, UL33 alters host cell signaling in a constitutive fashion (35–37), though our understanding of UL33-mediated signaling in epithelial cells and how this might impact infection remains unexplored. Thus, to begin to understand the basic biological functions of UL33 and gain insight its requirement for efficient epithelial cell replication, we created stable ARPE-19 cell lines containing doxycycline (DOX)-inducible expression constructs for each of the four HCMV vGPCRs. We used the pSLIK-Venus lentiviral backbone (57), which also allows us to sort transduced cells via flow cytometry, since the Venus fluorescent protein is constitutively expressed. Moreover, the vGPCRs were FLAG-tagged at the C-terminus to facilitate analyses of protein expression. Following lentiviral transduction of the ARPE-19 cells, Venus positivity was between 25 and 50% for each of the four vGPCR transduced lines. Venus-positive cells were sorted by flow-based cell sorting, resulting in stable lines with 85% to 95% Venus positivity. Using a standard immunoprecipitation/western approach, we demonstrate that each of the vGPCR proteins is expressed after DOX induction (**Fig. 8**). Moreover, this DOX inducible system is tightly regulated, as the protein abundance for each vGPCR in the absence of DOX is below the level of detection for the assay.

**Figure 8.**
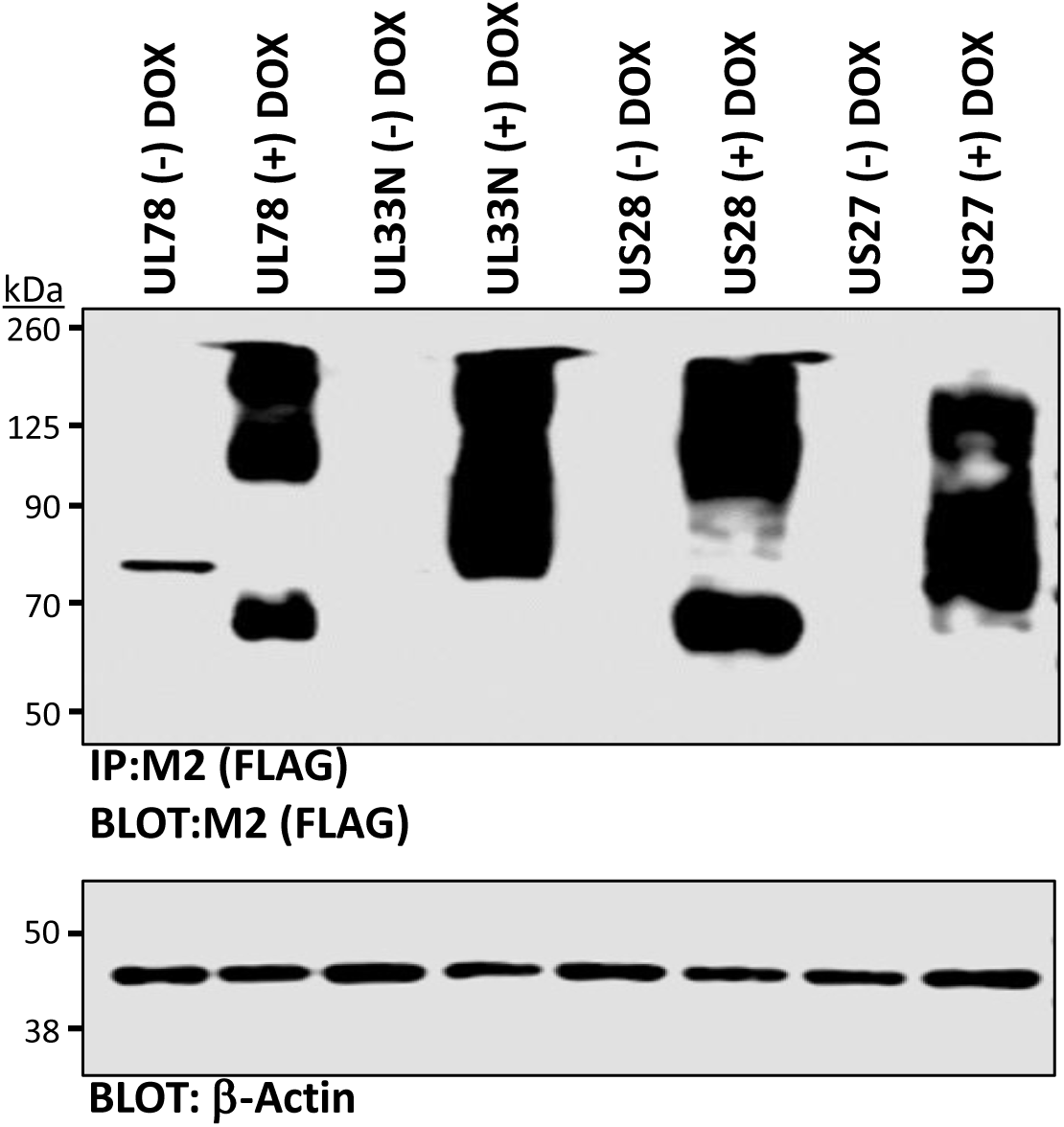
Stably transduced ARPE-19 cells support tightly regulated, doxycycline-dependent vGPCR protein expression. ARPE-19 cells were transduced with the pSLIK-Venus lentivirus expressing one of the four vGPCRs (UL33N [full-length], UL78, US27, or US28) under the control of a doxycycline (DOX)-inducible promoter and with a C-terminal FLAG epitope tag. Once stably transduced ARPE-19 cells were selected via the Venus tag, the cells transduced with pSLIK-Venus-UL78, pSLIK-Venus-UL33N, pSLIK-Venus-US27, or pSLIK-Venus-US28 were treated +/- DOX (1.0 *µ*g/mL) for 48 h. Cells were harvested, lysed in RIPA buffer, and vGPCR proteins were immunoprecipitated using anti-FLAG M2 agarose beads. Samples were then probed for the FLAG epitope tag (to detect each vGPCR) or β-actin as a loading control. The blots shown are representative of three independent experiments.

Once we confirmed expression of each vGPCR-encoded protein in the stable ARPE-19 cells (**Fig. 8**), we next examined the signaling activities of these vGPCRs. Prior work reveals US28 constitutively activates Gαq and PLC-β leading to IP3 accumulation (58), but much less is known about the other three vGPCRs UL33, UL78, and US27. The positional homolog of UL33 in murine cytomegalovirus (MCMV), called M33, is a strong inducer of the Gαq/PLC-β signaling (51, 59); however, published data with HCMV UL33 remains controversial, with some studies showing marginal Gαq/PLC-β signaling and others showing a complete lack of Gαq/PLC-β signaling (35–37). Moreover, similar to the MCMV M33 gene, UL33 is expressed as a long splice variant, giving rise to a 412 amino acid protein (UL33N), or a shorter splice variant, giving rise to an amino terminally truncated 390 amino acid protein (UL33S) (60). To begin to understand UL33-mediated functions in epithelial cells, we used the full length UL33N protein for these initial experiments. To assess Gαq/PLC-β signaling, we induced expression of each of the vGPCRs with DOX for 48 h, radiolabeled cells with [^3^H]-myoinositol, and ran standard assays to determine PLC-β activity by measuring inositol phosphate (IP) accumulation (50). Confirming earlier studies from our labs and others (50, 58, 59, 61), we observed strong induction of PLC-β activity from the US28 vGPCR (**Fig. 9A**). Interestingly, we also observed a strong induction of PLC-β activity when we induced UL33N expression. This is in contrast to US27 and UL78, neither of which were capable of inducing Gαq/PLC-β signaling (**Fig. 9A**). To compare the levels of signaling induced by the US28 and UL33N vGPCRs, we then performed a dose response assay with increasing concentrations of DOX. Both US28 and UL33N induced significant PLC-β signaling over a dose response curve from 10 ng/ml to 1 µg/ml DOX, and the level of signaling between these two vGPCRs is relatively comparable, although UL33N is slightly weaker in its ability to activate this signaling pathway in the ARPE-19 cells (**Fig. 9B**). Nonetheless, these data indicate that US28 and the full-length UL33 protein, UL33N, both induce PLC-β signaling in ARPE-19 epithelial cells.

**Figure 9.**
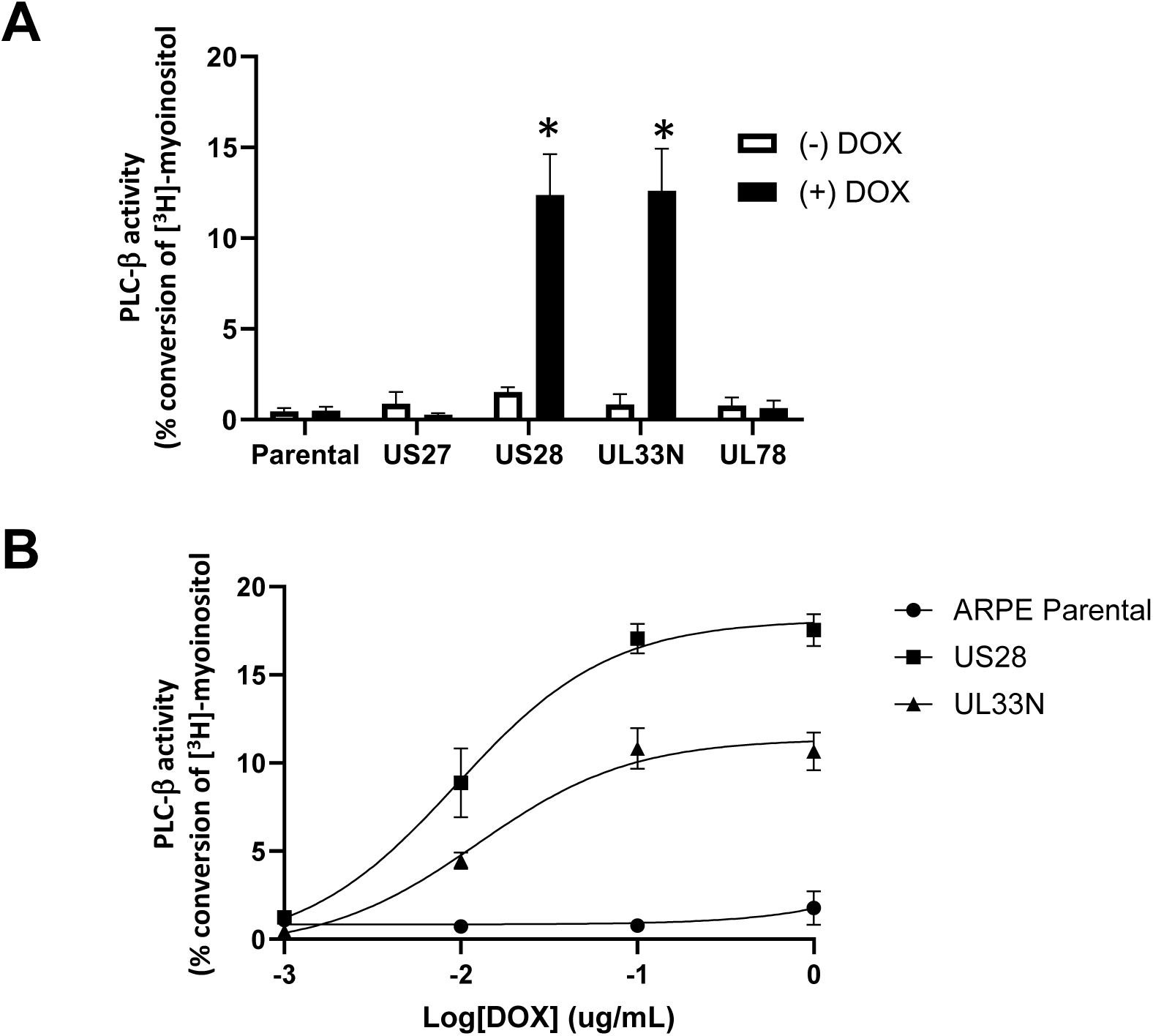
Both US28 and UL33N induce PLC-β activity in epithelial cells. (**A**) ARPE-19 cells stably expressing the indicated vGPCR were maintained for 48 h in the absence or presence of doxycycline (DOX; 1.0 *µ*g/mL), after which ^3^H-myoinositol was added to a final concentration of 1.0 *µ*Ci/well. Cells were incubated overnight in the ^3^H-myoinositol-containing media before the PLC-β activity assay was performed using a standard assay for detection of total inositol phosphates. As a control, non-transduced parental ARPE-19 cells (Parental) were included. (**B**) The assay described in (**A**) was performed as described, except with increasing doses (0, 0.01, 0.1, 1.0 *µ*g/ml) of DOX prior to measuring PLC-β activity in stably transduced ARPE-19 cells expressing US28 or UL33N, as well as the non-transduced ARPE Parental line. (**A,B**) Averages are shown with SEM error bars from three independent experiments performed in triplicate. Pairwise t tests between the induced and uninduced groups were used to determine statistical significance; \**p* < 0.05.

Although Gαq is typically the G-protein responsible for PLC-β activation and IP accumulation, Gαi can also activate PLC-β through its Gβγ subunits (62–64). Therefore, to determine if UL33N is coupled to Gαq, we employed a pharmacological approach. We previously showed US28 is coupled to Gαq using the pharmacological inhibitor, YM254890, results which we confirmed using Gαq-deficient mouse embryonic fibroblasts (53, 54). Therefore, we examined UL33N signaling through PLC-β using YM254890. Corroborating earlier studies, we found that US28 signaling was inhibited by YM254890, and interestingly, we also observed that UL33N was similarly inhibited by the compound (**Fig. 10**). These results confirm that UL33N is similarly coupled to Gαq, leading to PLC-β activation and IP accumulation. As discussed above, the *UL33* gene can give rise to a variant protein deleted for the amino terminal 22 amino acids, termed UL33S. Previous studies from our lab using the MCMV M33 vGPCR indicated that the M33 amino terminus is required for proper maturation, and an amino truncated M33 protein was incapable of inducing Gαq/PLC-β signaling (51). We therefore investigated if the amino terminus of UL33 was similarly required for Gαq/PLC-β signaling. To this end, we transduced ARPE-19 cells with a lentiviral expression vector and created a stable cell line expressing UL33S. We evaluated protein abundance by immunoprecipitation/western blot, which indicated that UL33S is expressed at levels similar to or greater than that of UL33N (**Fig. 11A**). We then assessed PLC-β activity as above, and in contrast to the full length 412 amino acid UL33N protein, the N-terminal truncated 390 amino acid UL33S protein is defective in stimulating PLC-β signaling (**Fig. 11B**), indicating region(s) within the amino terminus of UL33 are important for driving PLC-β signaling in ARPE-19 cells.

**Figure 10.**
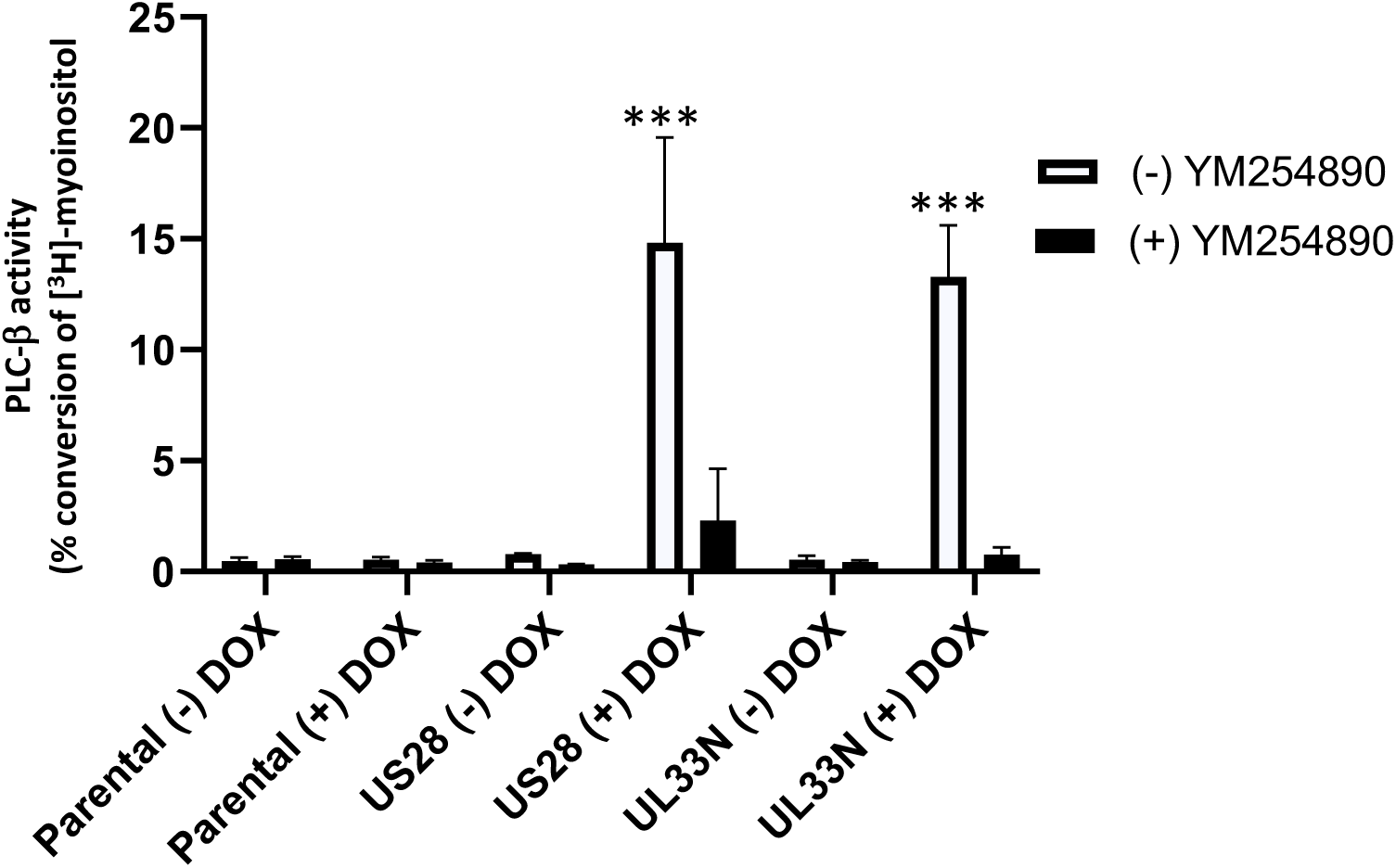
Both US28 and UL33N signal through Gαq to induce PLC-β activity in epithelial cells. ARPE-19-pSLIK-Venus-US28 or ARPE-19-pSLIK-Venus-UL33N were maintained for 48 h +/- doxycycline (DOX;1.0 *µ*g/mL), after which ^3^H-myoinositol was added to a final concentration of 1.0 *µ*Ci/well. Cells were incubated overnight in the ^3^H-myoinositol-containing media before the Gαq inhibitor, YM254890 (100 nM), was added. After allowing the inositol phosphates (IPs) to accumulate for 2 h, the total IPs were detected to determine PLC-β activity. Averages are shown with SEM error bars from three independent experiments, each performed in triplicate. 2-way ANOVA was used to determine statistical significance; \*\*\*\**p* < 0.0001.

**Figure 11.**
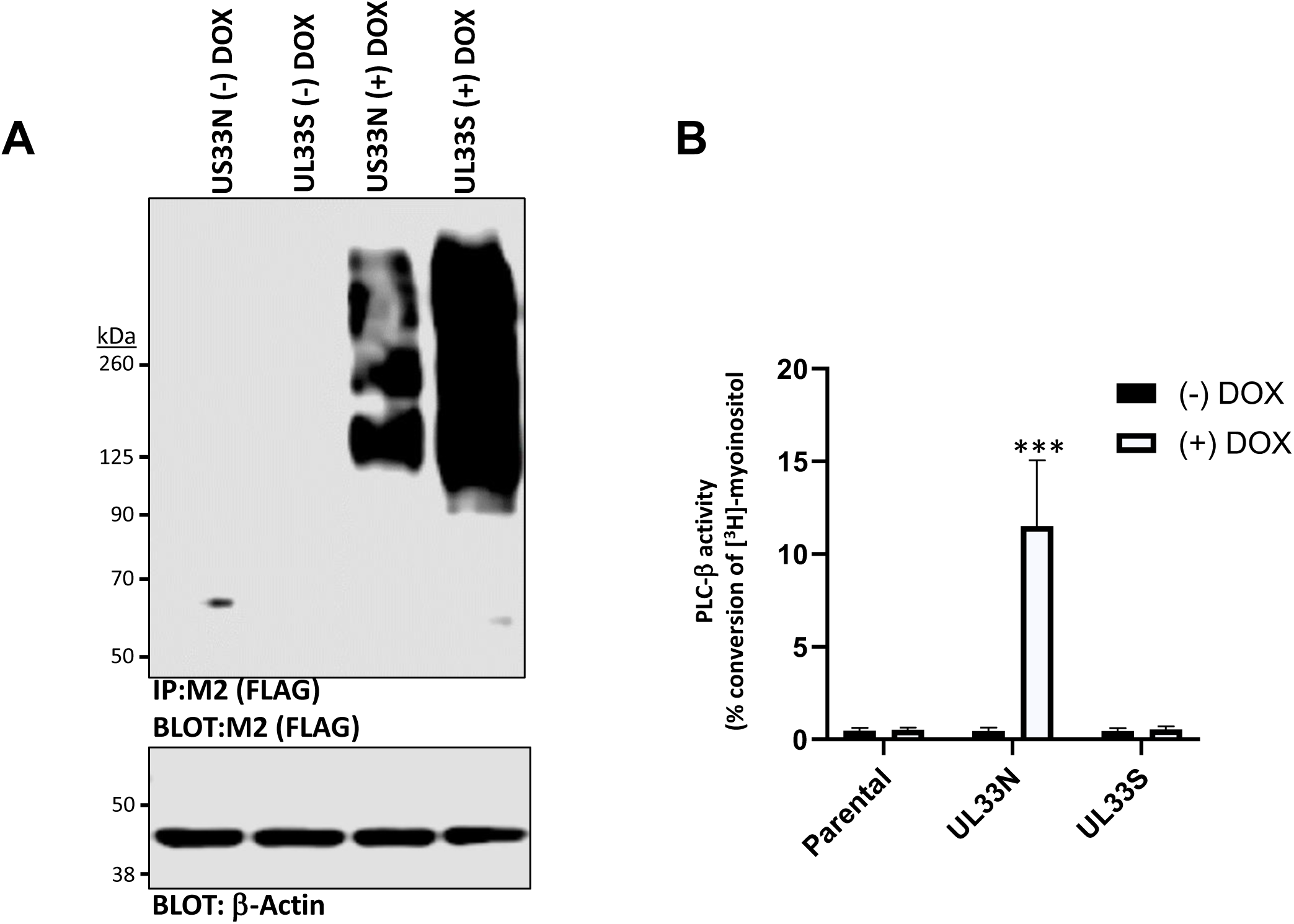
PLC-β activity is induced by UL33N, but not UL33S, in epithelial cells. Stable ARPE-19 cell lines were constructed using two splice variants of UL33. “UL33N” contains an additional Exon encoding 22 amino acids on the N-terminus, which is absent in “UL33S.” (**A**) Immunoprecipitation - western blot analysis was performed as described in Fig. 2 with protein extracts from the UL33N and US33S stable ARPE-19 cells following +/- doxycycline (DOX; 1.0 *µ*g/mL) for 24 h. The blot shown is representative of three independent experiments. (**B**) As described in Fig. 4, inositol phosphate (IP) accumulation was assessed to determine PLC-β activity in response to UL33N or UL33S proteins. Averages are shown with SEM error bars from three independent experiments performed in triplicate. Pairwise t tests between the induced and uninduced groups were used to determine statistical significance; \*\*\*\**p* < 0.0001.

### *UL33* has unique 5’ UTRs and encodes additional protein variants

While published gene expression and genome annotation studies indicate that UL33 can exist as either 412 amino acid UL33N or 390 amino acid UL33S variants (60), our transcriptomic data indicates that transcription within the *UL33* gene region is more complex. Our polysome pulldown/RNAseq data reveals additional transcriptional start sites for *UL33*, which can generate mRNAs with additional upstream exons and distinct five prime untranslated regions (5’ UTRs) (**Supplemental Table 1**). These data indicate there are four different exons (Exons 1, 2, 3, and 4) that can be used to form mRNAs expressed from the *UL33* gene region, rather than the two (Exons 3 and 4) originally reported to give rise to the previously discussed UL33N and UL33S proteins, respectively (**Fig. 12A, Supplemental Table 1**). The additional two exons (now called Exons 1 and 2), were both located upstream of the original exons (now called Exons 3 and 4), meaning that these exons could be independently utilized as distinct 5’ UTRs. We also noted that there was a third methionine start site upstream of the “UL33N” start site in transcripts containing extended versions of Exon 3. This could code for an additional variant UL33 protein, which is 24 amino acids longer than what was originally considered the full-length version of UL33N (**Fig. 12B**). Given the additional complexity within the *UL33* coding region, we designed and built new expression constructs for each of these variant *UL33* transcripts so that we could determine 1) if the unique 5’ UTRs affected translational efficiency, as measured by protein expression and signaling activity, and 2) if the novel protein product encoded by the extended Exon 3 mRNA, adding 24 amino acids to the N-terminus, was capable of activating Gαq/PLC-β signaling. We then created stable ARPE-19 cells expressing each of the *UL33* gene transcripts, as described above. We found that all four of our new *UL33* gene constructs led to the production of stable proteins (**Fig. 13A**). While the UL33 E2 and UL33 E3+54-containing constructs resulted in somewhat increased levels of UL33 expression (**Fig. 13A**), quantification of protein abundance from multiple, independent experiments did not indicate significant differences in the abundance of the four new constructs (data not shown). Therefore the variant 5’ UTRs found in Exon 1 (E1)- and Exon 2 (E2)-containing mRNA do not dramatically impact translation efficiency outside the context of infection in ARPE-19 cells (65). Moreover, when we tested these variants for their ability to induce Gαq/PLC-β signaling, we found that all the variant transcripts resulted in UL33 proteins capable of stimulating equivalent levels of IP3 accumulation (**Fig. 13B**). Taken together, these data suggest *UL33* expression is more complex than originally believed, leading to unique 5’UTR containing transcripts and previously undescribed protein variants. Further, the variant 5’UTR sequences and the additional protein variant encoded by exon E3+201-containing transcripts are unlikely to dramatically affect UL33-dependent signaling in infected cells.

**Figure 12.**
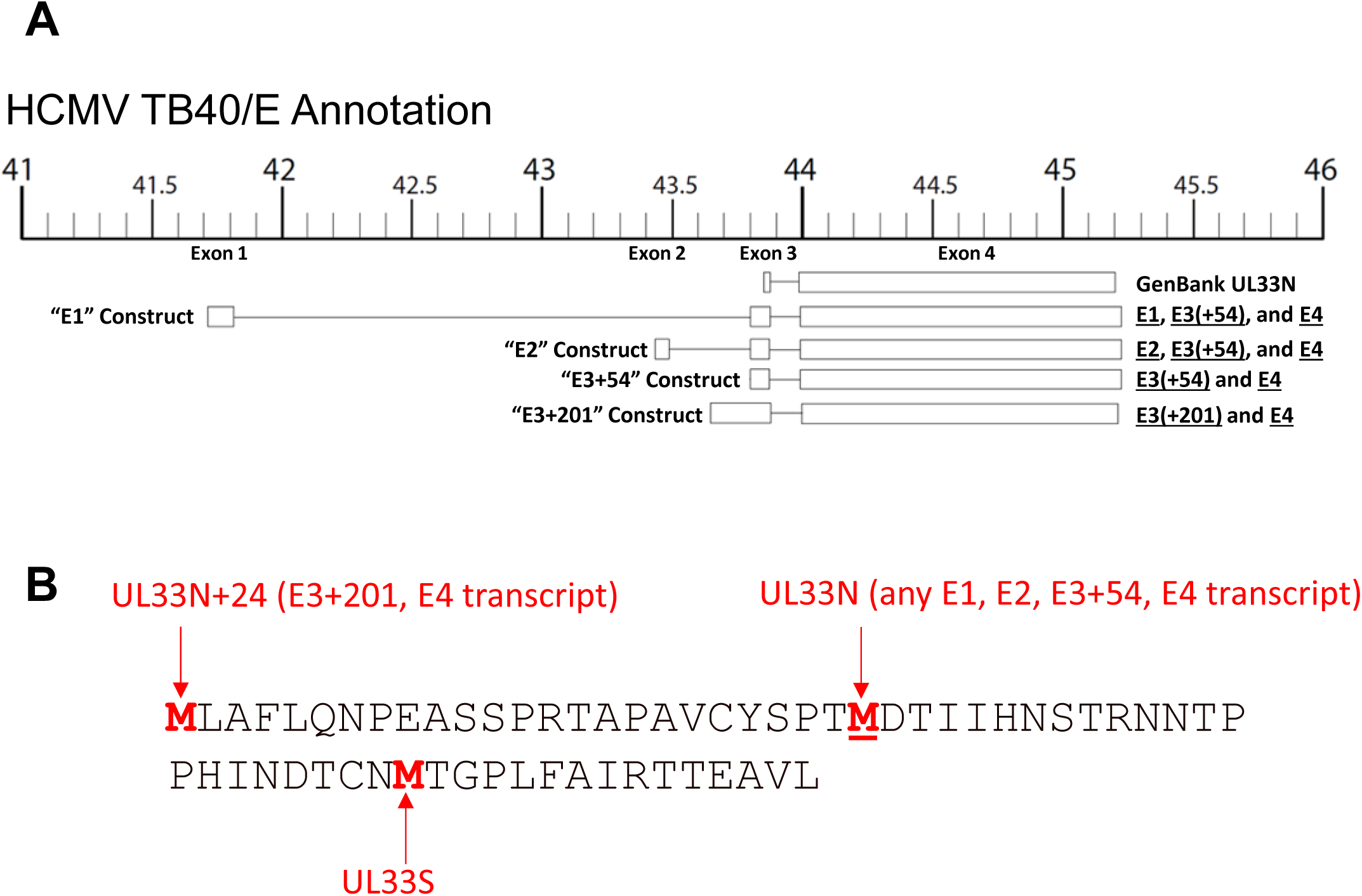
RNAseq analyses from polysome fractions indicate the potential for additional RNAs, which once translated, could result in UL33 protein variants. (**A**) Schematic of the four potential exons that combine to form UL33 mRNAs identified from Polysome/RNAseq data (Supplementary Table 1). Numbers represent TB40/E coordinates, and the Exons are labeled below the scale. UL33 expression constructs were created as depicted, representing each of the different, potentially-encoded UL33 mRNAs. Transcripts containing Exon 3 could be two different lengths (indicated by E3+54 or E3+201), with the longer transcript (E3+201) containing the coding potential to produce a novel UL33 protein variant with 24 extra amino acids on the amino terminus. (**B**) Amino acid sequence of the amino terminal region of UL33 depicting the three possible start sites (red methionine, M), UL33N+24, UL33N, and UL33S.

**Figure 13.**
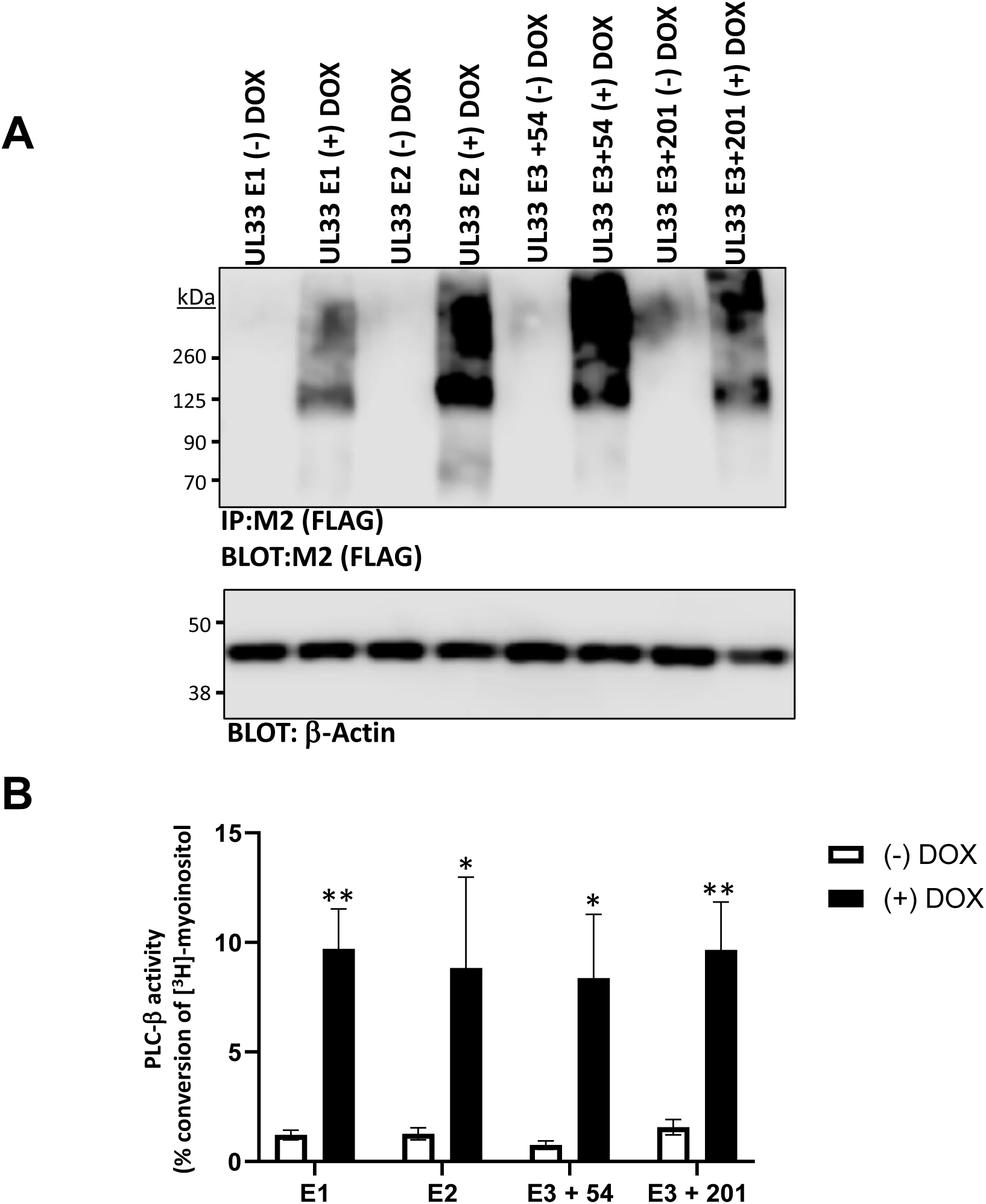
UL33 variants produce stable proteins that increase PLC-β activity in epithelial cells. (**A**) ARPE-19 cells were stably transduced with lentiviral constructs expressing each of the various UL33 gene product variants under the control of a doxycycline (DOX)-inducible promoter. Each stable cell line was then assessed for UL33 variant protein abundance by immunoprecipitation - western blot, as described in Fig. 2. The blots shown are representative of three independent experiments. (**B**) The accumulation of inositol phosphates (IPs) was assessed as a measure of PLC-β activity, as described in Fig. 4. Averages are shown with SEM error bars from three independent experiments, each performed in triplicate. Pairwise t tests between the induced and uninduced groups were used to determine statistical significance; \**p* < 0.05, \*\**p* < 0.01.

## Discussion

Though precise functions for each of the HCMV vGPCRs are not well-characterized, a growing body of evidence indicates the vGPCRs play a variety of roles in HCMV pathogenesis, including those central to the establishment and maintenance of latency (24, 30, 54, 66–75), as well as reactivation (29, 39). Here, using a virus in which we deleted the ORFs for each vGPCR (TB40/E-Δall), we show these vGPCRs are required for efficient replication in primary human salivary gland epithelial cells (hSGEs), as well as the model epithelial cell line, ARPE-19 (**Fig. 1**). Given the central role for the HCMV vGPCRs in promoting replication in primary salivary cells and similar results in the ARPE-19 cells, we posited that utilizing ARPE-19 cells were a valid model to investigate the functional biochemical properties of the vGPCRs in epithelial cells. We found all four vGPCR proteins are expressed during infection (**Fig. 2**), consistent with prior work examining UL78 (27) and US28 (48, 49). As little is known about UL33 and how it contributes to efficient replication in epithelial cells, we focused on UL33 herein. Our findings reveal deletion of the *UL33* ORF resulted in a virus that displayed attenuated viral replication in ARPE-19 epithelial cells (**Fig. 5**) and was severely crippled in its ability to promote all three phases of the lytic cascade, measured at both the mRNA and protein levels (**Figs. 6-7**). As UL33 signals constitutively in certain cell types (35–37), we hypothesized UL33-mediated signaling may contribute to its ability to promote efficient viral growth in epithelial cells. To begin to address this, we evaluated UL33 constitutive activation of PLC-β and found the full-length UL33 (UL33N), but not the truncated form (UL33S), couples to Gαq to potentiate PLC-β activation (**Figs. 9-11**). Our data also reveal there are four different UL33 exons, which are actively incorporated into mRNAs, thus creating potentially novel 5’UTRs for these UL33 mRNAs (**Fig. 12**). However, each of the four UL33 proteins translated from variant transcripts were abundantly expressed and constitutively activate Gαq/PLC-β (**Fig. 13**). Taken together, our findings reveal that the vGPCRs contribute to successful replication in epithelial cells, due at least in part, to the expression of UL33.

Initially thought of as dispensable for lytic replication (76, 77), continually emerging data support important biological functions for the vGPCRs during all phases of infection. Perhaps it is unsurprising that UL33 is important for HCMV infection, as its rodent counterparts, MCMV M33 and RCMV R33, play important roles in facilitating replication *in vivo* in epithelial cells, like those comprising the salivary epithelium (31–34). Our study indicates that deletion of UL33 from the HCMV genome is at least partially responsible for the effect we observe when all four vGPCRs are deleted, indicating that like its rodent counterparts, HCMV UL33 is also important for viral replication. However, it is also plausible that the other vGPCRs function at different stages of epithelial cell infection. Indeed, UL78 is required for efficient HCMV entry into epithelial cells (27). In further support of this hypothesis, we observed differential kinetics of protein accumulation for the vGPCRs (**Fig. 2**). While we only focused on two time points herein, we are keen to further elucidate the expression patterns of these vGPCR proteins during infection, especially in hSGE cells. Since the hSGE cells support low-level, persistent lytic HCMV replication for four to five weeks post-infection (13), it will be interesting to examine vGPCR expression at these very late time points post-infection and determine if the vGPCRs play a role in supporting this sustained, persistent lytic infection.

Our data reveal that US28 and UL33 both robustly activated PLC-β in ARPE-19 epithelial cells, as measured by inositol phosphate accumulation. Though US28 constitutive activation of Gαq-coupled PLC-β activity was previously shown in multiple cell types (46, 49, 58, 61, 78), we were intrigued with the finding that UL33 constitutively activated high levels of PLC-β activity in our experiments. This is somewhat distinct from other published data showing UL33 weakly activates PLC-β activity (35–37). A multitude of reasons, however, could account for such differences. For example, we utilized stable transduction for our cell lines rather than transient transfections. Additionally, the cell type used likely impacts results, as is the case with US28 (46). Much of the published data on UL33 has come from transformed cell lines, such as HEK-293T (35) or COS-7 (36), which do not support HCMV infection. In our studies, we utilized the epithelial cell line, ARPE-19, which is permissive to HCMV infection and may reflect a more biologically relevant cell type in which to investigate UL33 signaling. We also plan to create and analyze signaling in stable, vGPCR-expressing hSGE cells, representing perhaps the most clinically relevant model to interrogate such questions in tissue culture. Nonetheless, we were intrigued by the ability of UL33 to activate PLC-β and confirmed that this signal was a result of Gαq coupling using the Gaq inhibitor, YM-254890. It would seem somewhat redundant for HCMV to encode for two separate vGPCRs, UL33 and US28, which signal through the same heterotrimeric G protein. However, there is some difference in the expression kinetics between US28 and UL33 in HCMV infected epithelial cells (**Fig. 2**), providing potential insight into the apparent redundancy. The apparent US28- and UL33-mediated redundancy in activating Gαq/PLC-β signaling may reflect the need for the virus to maintain high levels of this signaling arm at different times post-infection. Future experiments aimed at analyzing UL33-dependent signaling in infected cells at various times post-infection, as well as during low-level hSGE persistent infection, will provide additional insights into this point. On the other hand, it is not surprising that HCMV has evolved to target Gαq/PLC-β via redundant mechanisms, as this is true for other cellular targets of HCMV (e.g., HCMV US2-, US3-, US26-, and US11-mediated downregulation of MHC Class I (79–85).

It is important to note that we only investigated constitutive signaling in these assays; it may prove worthwhile to further investigate possible ligand-induced signaling from each of the receptors (US28 and UL33), as this could provide a greater understanding as to why the HCMV genome would encode for two receptors which signal through the same G protein. Previously published work revealed that US28 is activated by various human CC-chemokines (RANTES, MCP-1, MCP-3, and MIP-1α), as well as fractalkine, a human CXC3-chemokine (58). However, there is currently no evidence for UL33 ligand-induced signaling, and thus, UL33 remains an ‘orphan receptor’. Moreover, our examination of Gαq signaling using the UL33 variant that encodes an amino terminal truncated protein (UL33S) revealed that the N-terminus is pertinent to the protein’s ability to signal through PLC-β in epithelial cells. There are two potential reasons for the failure of the truncated UL33S variant to signal though Gαq. First, it is possible that losing the N-terminus of the protein prevents the protein from trafficking to the cell surface, in turn blunting its ability to signal since it is not in the correct subcellular location. Second, the N-terminal domains of some GPCRs can play a significant role in activating the receptors through the utilization of peptide sequences in the amino terminus that function as “tethered ligands”. Both protease activated receptors (PARs) and members of the adhesion family of GPCRs use this mechanism to promote signaling (86–88). While we have only investigated constitutive signaling from UL33 proteins, it is possible that the N-terminus could act as a ligand for self-activation of the receptor. We will actively pursue this possibility in future experiments.

Our data also reveal two new, previously undescribed exons, which can potentially serve as novel 5’ UTRs for UL33 mRNA. Interestingly, while the MCMV UL33 ortholog, M33, exists as a two-exon transcript (60), the guinea pig CMV (GPCMV) UL33 ortholog, GP33, is expressed as a three-exon transcript with an additional exon located upstream (89). We now demonstrate that HCMV UL33 gene expression is more complex and similar to that of GPCMV GP33. We found four different UL33 exons, which are actively incorporated into mRNAs, thus creating potentially novel 5’UTRs for the UL33 mRNAs. This is intriguing, as the HCMV TB40/E annotation in GenBank (KF297339.1) describes only two exons. The GenBank annotation describes the upstream exon (which we now refer to as “Exon 3”) as having 24 base pairs; however, our polysome/RNAseq data shows two distinct lengths of Exon 3 (E3), with either an additional 54 or 201 base pairs. The addition of 201 base pairs to Exon 3 creates a novel UL33 protein, with an additional 24 amino acids on the amino terminus of the UL33N open reading frame. Our findings indicate that each of the four UL33 proteins made from variant transcripts were efficiently expressed and able to constitutively activate PLC-β. Moreover, we found that the UL33 protein with the extended amino terminus was equally competent for activation of Gαq/PLC-β signaling. Additionally, while the UL33 amino terminus is important for signaling, these findings indicate that the addition of 24 amino terminal amino acids does not impact its function. Thus, UL33 has multiple options for 5’UTR utilization to make stable UL33 proteins that signal through Gαq. What is the relevance of the two additional transcriptional start sites that give rise to transcripts that synthesize functional UL33 variants? At present, the answer is unclear, but it is attractive to posit this indeed may prove important, as the promoters upstream of these start sites might be differentially regulated thus altering the transcript levels in certain cell types or during specific phases of infection.

Taken together, this study provides new clues into the epithelial cell functions of HCMV vGPCRs and UL33, in particular. Future experiments aimed at linking the biochemical functions of UL33, including its ability to potentiate signaling, with its biological ability to drive viral replication in epithelial cells will provide important insights into how HCMV can persist in the salivary epithelium and promote horizontal dissemination.

## Materials and Methods

### Cell line maintenance and isolation of primary salivary cells

293T, ARPE-19, and HS68 human foreskin fibroblast (HFF) cells were acquired from the American Type Culture Collection (ATCC). Primary newborn human foreskin fibroblasts (NuFF-1) were obtained from GlobalStem. 293T are epithelial-like cells isolated from a human embryonic kidney and were maintained in Dulbecco’s modified Eagle’s medium (DMEM) (Corning) supplemented with 10% fetal bovine serum (FBS; Corning) and 1% penicillin-streptomycin (P/S). ARPE-19 cells are adult human retinal pigment epithelial cells and were maintained in DMEM–F-12 medium (50:50; Corning) with 10% FBS, 1% P/S, and in some cases, 1% amphotericin B. HS68 HFF cells were maintained in DMEM supplemented with 10% FetalClone III serum (HyClone) and 1% P/S. NuFF-1 cells were maintained in DMEM, containing 10% FBS (MilliporeSigma), 2 mM L-glutamine, 10 mM HEPES, 0.1 mM non-essential amino acids, and 100 U/ml each of penicillin and streptomycin.

Primary human submandibular and parotid salivary gland-derived epithelial (hSGE) cells originated from patient salivary tissue at the University of Cincinnati’s Department of Otolaryngology; these cells were maintained in BEGM growth medium (BEBM media containing the BEGM bullet kit; Lonza) and supplemented with 4% charcoal-stripped FBS (Gibco). This study was reviewed and approved by the Institutional Review Board at the University of Cincinnati (Federal-wide assurance number 00003152, IRB protocol 2016-4183). All cells were maintained at 37°C with 5% CO2.

### Recombinant viruses, viral stock generation, and titers

The viruses used in this study were derived from the parental, clinical, bacterial artificial chromosome (BAC)-derived HCMV isolate, TB40/E-BAC (clone 4) (90). TB40/E*mCherry* (WT) (45), TB40/E*mCherry*-UL33-3xF (UL33-3xF) (39), TB40/E*mCherry*-UL78-3xF (UL78-3xF) (27), TB40/E*mCherry*-US27-3xF (US27-3xF) (45), TB40/E*mCherry*-US28-3xF (US28-3xF) (46), TB40/E*mCherry*-allΔ (allΔ) (46), TB40/E*mCherry*-UL33Δ1 (UL33Δ1) (39), and TB40/E*mCherry*-UL33Δ2 (UL33Δ2) (39) generation and characterization were previously described in the indicated reference. Virus stocks were generated as previously described (91). In brief, virus genome-containing BAC DNA was isolated and purified, after which DNA was electroporated into NuFF-1 or HS68 fibroblasts to generate a p0 stock. This stock was then used for expansion to generate a p1 stock on naïve fibroblasts, which, following 100% cytopathic effect (CPE), was harvested. For experiments in Figs. 3-7, infectious supernatant was purified through a 20% sorbitol cushion containing 0.1M Tris, pH 7.2 and 2.0 mM MgCl2, resuspended in X-VIVO 15 media (Lonza) and 3% BSA (1:1 ratio), flash-frozen in liquid nitrogen, and then stored at -80C. Viral stocks were titered by 50% tissue culture infectious dose (TCID50) assay on naïve fibroblasts or by quantitation of IE positivity on naïve fibroblasts in flow cytometry assays (IU).

To analyze the ability of TB40/E-Δall to infect epithelial cells relative to that of fibroblasts, hSGE or ARPE-19 cells were infected at an MOI of 2 IU/cell. Concurrently, an equal number of HS68 HFF cells were infected at the same MOI. After two days of infection, cells were harvested and fixed in 70% ethanol (EtOH) for at least 30 minutes (min) at 4°C. Samples were then centrifuged at 800 × *g* for five min before being resuspended in 500 μL permeabilization buffer (1X DPBS containing 0.5% BSA and 0.5% Tween-20) and incubated at 4°C for 10 min. Cells were pelleted again, then resuspended in staining buffer (1X DPBS containing 0.5% BSA) and a 1:250 dilution of IE1 antibody conjugated to Alexa Fluor® 488 (EMD Millipore Corp. MAB810X). After incubating for 1-2 hours in the dark at room temperature, cells were analyzed by flow cytometry for the proportion of cells which were IE1 (Alexa Fluor® 488) positive. Infectivity Indexes were calculated by taking the percent of IE1 positive epithelial cells and dividing by the percent IE1 positive HS68 HFF cells from the same experiment. To determine the viral titers produced from the primary hSGE cells, cells derived from two independent submandibular glands and two independent parotid glands were infected at an MOI of 2 IU/cell. Supernatants from these cells were harvested at the indicated days post-infection (dpi) and flash frozen. Once the entire time course was harvested, supernatants were thawed and tittered on HS68 HFF cells, as described previously (43).

### Lentiviral transductions of ARPE-19 cells to create stable cells expressing vGPCRs

Viral G protein-coupled receptor (vGPCR) gene sequences were obtained from the AD169 viral genome and synthesized by GenScript. Synthesized vGPCRs were inserted into puc-57 plasmids incorporating a Kozak sequence upstream of the translational start site and a FLAG-tag (DYKDDDDK) on the C-terminus. The vGPCR genes were then inserted into the donor pEN-TmiRC3 plasmid digested with SpeI and XbaI (57). pENTR-vGPCR plasmids were confirmed through Sanger sequencing, and the vGPCR gene sequences were recombined into the pSLIK-Venus (57) utilizing LR recombination, according to the manufacturer’s protocol with LR Clonase (Invitrogen) and confirmed through Sanger Sequencing. Lentiviral particles were produced utilizing 293T cells. psPAX2 (1.5 μg), pVSVg (1.0 μg), and pSLIK-vGPCR Venus (3.5 μg) DNAs were combined with the Mirus TransIT®-LT1 Reagent (Mirus Bio) and added dropwise to 10 cm plates containing 2-3×10^6^ 293T cells plated the previous day. Lentivirus was harvested from these cells after 3 d and used to transduce ARPE-19 cells. After 2 d, ARPE-19 cells were analyzed by flow cytometry to determine the percentage of cells which express Venus. The cells were then sorted using a BD FACSAria II cell sorter in the Cincinnati Children’s Research Flow Cytometry Core Facility to obtain a >95% Venus-positive population.

### RNA and Protein Analyses

For all assays, fibroblast or epithelial cells were infected with the viruses at the multiplicities of infection (MOI) as indicated in the text and figures. Samples were collected for either RNA or protein analyses at the time points indicated in the text and figures. Where indicated, phosphonoacetic acid (PAA) was added to cultures (100 µg/ml) following virus adsorption, as previously described (27).

For RNA analysis, total RNA was isolated at times post-infection indicated in the text. RNA was isolated using the High Pure RNA Isolation kit (Roche), according to the manufacturer’s instructions. Equal RNA (1.0 µg) was then used to generate cDNA using TaqMan Reverse Transcription Reagents and random hexamer primers (both from Roche), according to the manufacturer’s protocol. Equal volume of cDNA was then used for qPCR using primers specific for *UL123* (forward, 5’-GCCTTCCCTAAGACCACCAAT-3’; and reverse, 5’-ATTTTCTGGGCATAAGCCATAATC-3’), *UL44* (forward, 5’-TACAACAGCGTGTCGTGCTCCG-3’; and reverse, 5’-GGCGTGAAAAACATGCGTATCAAC-3’), *UL99* (forward, 5’-TTCACAACGTCCACCCACC-3’; reverse, 5’-GTGTCCCATTCCCGACTCG-3’), and glyceraldehyde-3-phosphate dehydrogenase (*GAPDH*: forward, 5’-ACCCACTCCTCCACCTTTGAC-3’; and reverse, 5’-CTGTTGCTGTAGCCAAATTCGT-3’). Transcripts were quantified using a standard curve with tenfold serial dilutions of a BAC-standard that contains a sequence fragment of *GAPDH* (92). Viral transcripts were normalized to host-encoded *GAPDH* and plotted as arbitrary units (AU). Primer sets had similar efficiencies (>85%) and linear ranges of detection for the BAC standard (linear between 15 and 32 copies; r^2^>0.98 for all experiments). Samples were assessed in triplicate using the CFX Connect (Bio-Rad).

For protein analyses by western blot, cells were lysed in radioimmunoprecipitation assay (RIPA) buffer (1.0% NP-40, 1.0% sodium deoxycholate, 0.1% SDS, 0.15 M NaCl, 0.01 M NaPO4, pH 7.2, and 2 mM EDTA) on ice for 1 h; samples were vortexed every 15 min. Protein lysates were denatured in 2% SDS at 95°C for 10 min for all proteins except UL33, which was denatured at 42°C for 10 min. Equal concentrations of protein lysates were separated by SDS-PAGE and transferred by semi-dry transfer to Protran nitrocellulose (Amersham). Proteins were detected using the following antibodies: anti-FLAG M2 (MilliporeSigma), diluted 1:7,500; anti-IE1 (clone 1B12, diluted 1:50) (93); anti-ICP36 (Virusys; to detect pUL44), diluted 1:2,500; anti-pp28 (clone 10B4-29, diluted 1:100) (94); anti-actin-horseradish peroxidase (HRP; MilliporeSigma), diluted 1:20,000; goat-anti-mouse-HRP (Jackson ImmunoResearch Labs), diluted 1:10,000.

To analyze expression of each of the 4 HCMV vGPCRs in HCMV-infected epithelial cells, ARPE-19 cells were infected at an MOI of 10 IU/cell to ensure robust infection utilizing TB40/E vGPCR-3xF viruses. Cell lysates were then collected as described above at 48 and 96 hpi by harvesting cells in 1.0 ml of modified RIPA buffer (1.0% Triton X-100, 1.0% sodium deoxycholate, 0.1% SDS, 0.15 M NaCl, 0.01 M Tris pH 7.5, 5 mM EDTA, and Complete Mini Protease inhibitors) and briefly sonicated on ice. Whole cell lysates were prepared by mixing equal volume of protein lysate with 3× Laemelli sample buffer (50 μl each). The remainder of the protein lysate was immunoprecipitated overnight at 4°C using anti-FLAG M2 agarose beads (MilliporeSigma). Beads were washed three times with 1.0 ml RIPA buffer before being resuspended in 3× Laemelli sample buffer (50 μL). Immunoprecipitates were separated by SDS-PAGE and transferred to nitrocellulose as described above. Immunoprecipitates were probed with M2 conjugated to HRP (MilliporeSigma). All primary antibodies were diluted 1:1,000 in Tris-buffered saline with 0.1% Tween® 20 detergent (TBST).

To confirm that our stable ARPE-19 cells were producing vGPCR proteins, 1×10^6^ cells were plated in 10 cm plates in complete media with or without doxycycline (1.0 μg/mL) for 2 d. Cells were harvested, lysed in 1.0 ml of modified RIPA buffer, and briefly sonicated on ice. Whole cell lysates were prepared by mixing equal volume of protein lysate with 3× Laemelli sample buffer (50 μl each). The remainder of the protein lysate was immunoprecipitated, washed, and resuspended as described above. Whole cell extracts from stable cells were probed using antibodies against β-actin conjugated to horseradish peroxidase (HRP) (Cell Signaling Technology) and analyzed by western blot, as described above.

### Immunofluorescence Analysis of UL33 subcellular localization

To evaluate protein localization by immunofluorescence assay (IFA), fibroblasts were grown on glass coverslips and infected as indicated in the text. At the time points indicated in the text, cells were washed with 1X phosphate-buffered saline (PBS), fixed with 2.0% paraformaldehyde, and permeabilized with 0.1% Triton X-100. Slides were then blocked in 2.0% BSA in PBS containing 0.2% Tween at 4°C. For time course studies, fixed cells were maintained under these conditions until all time points were collected. Slides were then probed with anti-FLAG M2 (MilliporeSigma), diluted 1:500 in blocking buffer. Slides were then stained with Alexa 488-conjugated anti-mouse secondary antibody (Molecular Probes), diluted 1:1,000. Coverslips were then mounted onto slides with SlowFade Gold antifade reagent with DAPI (Life Technologies). Images were collected using a Leica TCS-SP8-AOBS confocal microscope.

### PLC-β activity assays in stable ARPE-19 epithelial cells

To elucidate the phospholipase C beta (PLC-β) activity of the various vGPCRs, 1×10^5^ stable ARPE-19 cells per well were plated in 12-well plates in complete media and grown for 48 h with or without doxycycline (1.0 μg/mL). Media was then removed and replenished with MEM containing ^3^H-myoinositol (PerkinElmer) at a final concentration of 1 μCi/well. Cells were incubated overnight (approximately 16 h) in the ^3^H-myoinositol-containing media before the PLC-β activity assay was performed using standard assays for detection of total inositol phosphates as described previously (50).Where indicated, the assay was repeated utilizing the Gαq inhibitor, YM254890, at a final concentration of 100nM during the inositol phosphate accumulation step, as it has been previously shown that this concentration is sufficient for inhibiting Gαq activity (54).

### Polysomal Profiling

Transcription through the UL33 gene in human fibroblasts infected for 6, 24, 48, or 72 hours with HCMV TB40/E at a MOI=3 was analyzed by next generation sequencing of polysome-associated RNA. Polysomes were resolved using linear sucrose gradients as described previously (95). Briefly, cells were treated with 100 μg/ml cycloheximide for 10 min at 37°C prior to harvest. Cell pellets were resuspended in polysome buffer (20 mM Tris-HCl [pH 7.4], 140 mM KCl, 5 mM MgCl2) containing 0.1% Triton X-100 and 10 mM dithiothreitol (DTT) and disrupted by 5 passes through a 27-gauge needle. Nuclei were removed by centrifugation for 5 min at 2,500 × *g*, followed by centrifugation for 10 min at 13,000 × *g* to remove insoluble debris. The clarified cytoplasmic lysate was layered onto linear 10 to 50% sucrose gradients prepared in polysome buffer and spun in an ultracentrifuge (Becton-Dickinson) for 2 h at 18,062 x *g* in an SW41 swinging bucket rotor without brake. The gradient was fractionated using a gradient fractionation system (Brandel) with continuous absorbance monitoring at an optical density at 254 nm (OD254). Gradient fractions were extracted with TRIzol (Thermo Fisher Scientific) and treated with Turbo DNase (Applied Biosystems), to isolate pure polysome associated RNA. Once purified, RNA libraries where produced using NEBNext Ultra II Directional RNA Library Prep Kit for Illumina following standard library preparation protocol. Libraries were sequenced on a HiSeq HS4000 Sequencer (Illumina).

## Acknowledgements

We would like to thank Anil Menon, Stephen Waggoner, Katherine Vest, and James Bridges for their insight and thoughtful discussions throughout the course of this work. This work was supported by National Institutes of Health Training Grant, T32ES007250 (awarded to MRF and MJB), American Heart Association Predoctoral Fellowship, 20PRE35200360 (awarded to MJB), as well as National Institutes of Health grants, R01AI121028 and R21DE026267 (awarded to WEM), R01AI153348 (awarded CMO’C), and R01AI103311 and R21AI123811 (to NJM). The funders had no role in study design, data collection and interpretation, or the decision to submit the work for publication.

---

**Supplemental Table 1.** Read Counts from RNAseq analysis indicating UL33 transcription containing additional upstream exons.

|  | Exon 1<br>(TB40E coordinates) | Exon 2<br>(TB40E coordinates) | Exon 3<br>(TB40E coordinates) | Exon 4<br>(TB40E coordinates) | Description | Read count<br>supporting this |  |
| --- | --- | --- | --- | --- | --- | --- | --- |
| UL33 Genbank Annotation |  |  | 43847-43870 | 43992-45206 | Contains E3,E4 |  |  |
| 6hr | 41716-41814 |  | 43793-43870 | 43871-45426 | Contains E1,E3,E4* | 330 |  |
| **6hr** | **41716-41814** |  | **43793-43870** | **43992-45426** | **Contains E1,E3,E4** | **465** | **#** |
| **6hr** |  | **43435-43494** | **43793-43870** | **43992-45426** | **Contains E2,E3,E4** | **1480** | **#** |
| 6hr |  |  | 43672-43870 | 43992-45426 | Contains E3,E4 | 2885 |  |
| 6hr |  | 43435-43494 | 43793-43870 | 43871-45246 | Contains E2,E3,E4* | 406 |  |
| 24hr | 41747-41814 |  | 43793-43870 | 43871-44835 | Contains E1,E3,E4* | 1245 |  |
| 24hr |  | 43441-43494 | 43793-43870 | 43871-44835 | Contains E2,E3,E4* | 11489 |  |
| 24hr |  | 43462-43494 | 43642-43870 | 43871-44835 | Contains E2,E3,E4* | 3794 |  |
| 48hr | 41777-41814 |  |  | 43992-45206 | Contains E1,E4 | 27366 |  |
| 48hr |  |  | 43653-43870 | 43871-44837 | Contains E3,E4* | 16134 |  |
| 72hr | 41712-41814 |  | 43793-43870 | 43871-45012 | Contains E1,E3,E4* | 11813 |  |
| 72hr |  |  | 43646-43870 | 43871-45012 | Contains E3,E4* | 54531 |  |
| **72hr** |  |  | **43646-43870** | **43992-45206** | **Contains E3,E4** | **15925** | **#** |

\* Indicates RNAs contain E3 and E4 as well as the intervening intron between E3 and E4

